# Stability of germline structural variation under dietary and ethanol challenges

**DOI:** 10.64898/2026.02.09.704746

**Authors:** Ivan Pokrovac, Željka Pezer

**Affiliations:** Division of Molecular Biology, Ruđer Bošković Institute, Zagreb, Croatia

**Keywords:** Structural variation, copy number variation, germline mutations, environmental exposure, optical genome mapping, sperm

## Abstract

Environmental and lifestyle factors can modify the sperm epigenome and influence offspring phenotype, yet whether such exposures directly affect structural variation in the male germline remains unclear. Structural variants (SVs) arise primarily through replication-associated mechanisms and represent a major source of genomic diversity and disease risk. Here, we tested whether three physiologically relevant exposures - a Western diet, a low-protein diet, and chronic ethanol consumption - alter germline SV profiles in mice. Using optical genome mapping, we analyzed paired sperm and kidney samples from individual animals, enabling within-mouse comparisons between germline and somatic tissues. Across all conditions, putative de novo SVs constituted ∼7% of detected variants, with comparable proportions in sperm and kidney. No significant differences were detected between experimental and control groups in SV count, size distribution, variant allele fraction, or zygosity in either tissue. A genome-wide scan identified a small number of candidate loci with nominal SV frequency differences in Western diet-exposed sperm, none of which remained significant after multiple-testing correction. Together, these results indicate that common dietary and alcohol-related exposures do not induce detectable global changes in germline structural variation, suggesting that any environmental effects on SV formation are at most subtle or highly localized.

## Introduction

Genetic variation arises from mutations generated by intrinsic processes such as DNA replication and repair errors, as well as from exposure to endogenous and environmental genotoxic agents. Although a low mutation rate is essential for evolution and adaptation, excessive mutagenesis can have deleterious consequences, including cancer and heritable genetic disease. Among all forms of genetic variation, structural variants (SVs) - including deletions, insertions, duplications, inversions, and translocations - account for the largest fraction of divergent base pairs between mammalian genomes. A typical human genome harbors tens of thousands of SVs, collectively affecting a substantial proportion of the genome and contributing significantly to phenotypic diversity, adaptation, and disease susceptibility [1–8]. Copy number variants (CNVs), a major subclass of SVs, are particularly important due to their high mutation rate relative to single-nucleotide variants and their potential to disrupt gene dosage and regulatory architecture [9]. De novo CNVs are a frequent cause of developmental disorders and are often inferred to arise in the germline, with a strong paternal bias observed for large structural rearrangements [9–11]. Despite their biological and clinical importance, the extent to which environmental factors modulate SV formation remains far less studied than that of the underlying point mutations.

Experimental and observational studies suggest that environmental stressors can increase the rate of structural mutations. Elevated CNV frequencies have been reported following exposure to ionizing radiation, heavy metals, air pollution, and replication-stress-inducing agents [12–17]. However, several studies have shown that even non-mutagenic, physiologically relevant environmental exposures, such as dietary composition or nutritional stress, can directly induce structural variation. The most compelling direct evidence comes from controlled experiments in *Drosophila melanogaster*, which demonstrated that increased dietary yeast concentration - a physiologically relevant manipulation emulating periods of dietary excess - causes ribosomal RNA gene (rDNA) copy number reduction and instability [18]. Critically, these diet-induced rDNA changes were not confined to somatic cells but also occurred in the germline, leading to transgenerational inheritance of reduced rDNA copy numbers. Complementary evidence comes from a study on yeast, which showed that prolonged starvation in *Saccharomyces cerevisiae* leads to large-scale chromosomal rearrangements occurring at rates far exceeding spontaneous mutation frequencies, which are maintained through subsequent generations [19]. The mechanisms likely involve activation of stress-response pathways and altered recombination dynamics. While yeast represents a unicellular eukaryote with different genome organization than animals, these findings suggest that nutrient availability can directly modulate structural variation rates through non-mutagenic mechanisms.

Mechanistically, many SVs are thought to arise through replication-dependent repair pathways, such as fork stalling and template switching (FoSTeS) or microhomology-mediated break-induced replication (MMBIR) [20, 21]. These processes are particularly sensitive to replication stress and are exacerbated by collisions between the replication and transcription machinery [22, 23]. Evidence from yeast and cultured mammalian cells demonstrates that transcription-associated replication stress can promote CNV formation, especially at highly transcribed or environmentally responsive loci [24–26]. In unicellular organisms, such environmentally stimulated copy number variation has been proposed as a mechanism of rapid adaptation, sometimes referred to as “directed mutagenesis” [26, 27]. While such mechanisms are generally considered unlikely to operate broadly in metazoans due to stringent genome maintenance and homeostatic buffering, all molecular components involved in transcription-replication conflicts are evolutionarily conserved [26, 28].

Male germ cells represent a unique cellular context in multicellular organisms: spermatogenesis involves extensive mitotic and meiotic divisions, rendering germ cells particularly vulnerable to replication stress. Indeed, male gametes are known to be highly sensitive to environmental perturbations, including diet, metabolic status, alcohol exposure, and other lifestyle factors [29, 30]. A substantial body of work demonstrates that paternal environmental exposures can reprogram the sperm epigenome, with consequences for offspring phenotype. Diet, obesity, alcohol consumption, stress, and exercise have all been shown to alter sperm epigenetic profiles in animal models, and in some cases to influence metabolic or behavioral traits in subsequent generations [30–37]. These findings raise the possibility that, by altering chromatin structure, environmental factors might also influence the germline genome itself by affecting the accessibility of genomic regions to recombination and repair machinery. Indeed, such a connection has been proposed in a study on rats, which suggested that environmentally induced epigenetic changes in sperm can promote copy number variations in subsequent generations [38]. However, this study used a reproductive toxin with genotoxic potential as an inducing agent, and direct evidence linking relatively mild, physiologically relevant environmental exposures to epigenetic instability and increased germline SV formation in mammals is still lacking.

Collectively, these studies challenge the view that structural variation arises solely through spontaneous mutational processes, revealing instead that genomes exhibit dynamic plasticity in response to environmental conditions. This has implications for understanding the origins of natural genomic variation and the potential for rapid evolutionary adaptation. However, critical gaps remain in our understanding of these phenomena in mammalian and human systems. While evolutionary genomic analyses suggest that various environmental challenges like shifts in diet, temperature, parasite diversity, availability of light and oxygen, may shape population CNV patterns in humans and non-human species [7, 39, 40], these studies are largely observational and cannot establish whether such exposures directly induce these CNVs or whether selection acts on spontaneous variants. If physiologically relevant environmental factors can induce SVs in humans, this raises questions about the health consequences of environmental exposures. CNVs are known to contribute to disease susceptibility, and environmentally induced CNVs could potentially increase risk for genomic disorders, cancer, or other conditions. The possibility of transgenerational inheritance of induced SVs suggests that environmental exposures could have multigenerational consequences through germline effects. This aligns with growing evidence for transgenerational epigenetic inheritance and raises the possibility that parental or ancestral environmental exposures could influence offspring genomic stability.

Most studies of de novo CNVs rely on parent-offspring comparisons using somatic tissues, which limits detection to variants transmitted to the next generation and obscures allelic and mosaic variation present in germ cells [41]. Direct analysis of sperm genomes offers a more sensitive approach to detecting de novo germline variants and distinguishing them from somatic variation. We recently established a protocol for preparing high-molecular-weight DNA from mouse sperm for optical genome mapping (OGM), enabling genome-wide, single-molecule analysis of germline structural variation [42]. Such advances in OGM now provide opportunities to directly investigate how environmental factors influence structural variation in the germline. In this study, we used OGM to examine structural variation in matched sperm and kidney samples from individual mice exposed to three environmentally relevant conditions: a Western diet, a low-protein diet, and chronic ethanol consumption. By comparing germline (sperm) and somatic (kidney) SV profiles within the same individuals, we aimed to quantify putative de novo SV formation and to test whether these environmental exposures alter SV count, size, variant allele fraction, or zygosity. We further explored whether specific genomic regions exhibit altered SV frequencies under dietary stress. Together, this work provides a systematic assessment of the extent to which common environmental factors influence germline structural variation and evaluates both the power and limitations of optical genome mapping for detecting subtle mutational effects in vivo.

## Results

### Data Filtering and Quality Assessment

To assess the quality of the initial data and the de novo genome assemblies, we analyzed how the key quality indicators of each sample compare with the recommendations by the Bionano Genomics guidelines (Supplementary Table S1). One sample from the control group in the ethanol consumption experiment (CAM0006335) and another sample from the experimental group in the low-protein diet experiment (CAM0005996) exhibited extremely low diploid genome map N50 values (1.4 and 1.7 Mbp, respectively). Due to insufficient assembly contiguity, both samples were excluded from all downstream analyses.

We next examined the number and lengths of contigs (consensus genome maps) per chromosome across all remaining samples (Fig. 1). Assemblies of the sex chromosomes were consistently fragmented, exhibiting both a higher number of contigs and markedly shorter contig lengths than autosomes. Because such reduced contiguity can compromise SV calling accuracy, sex chromosomes were excluded from all SV analyses.

**Fig. 1.**
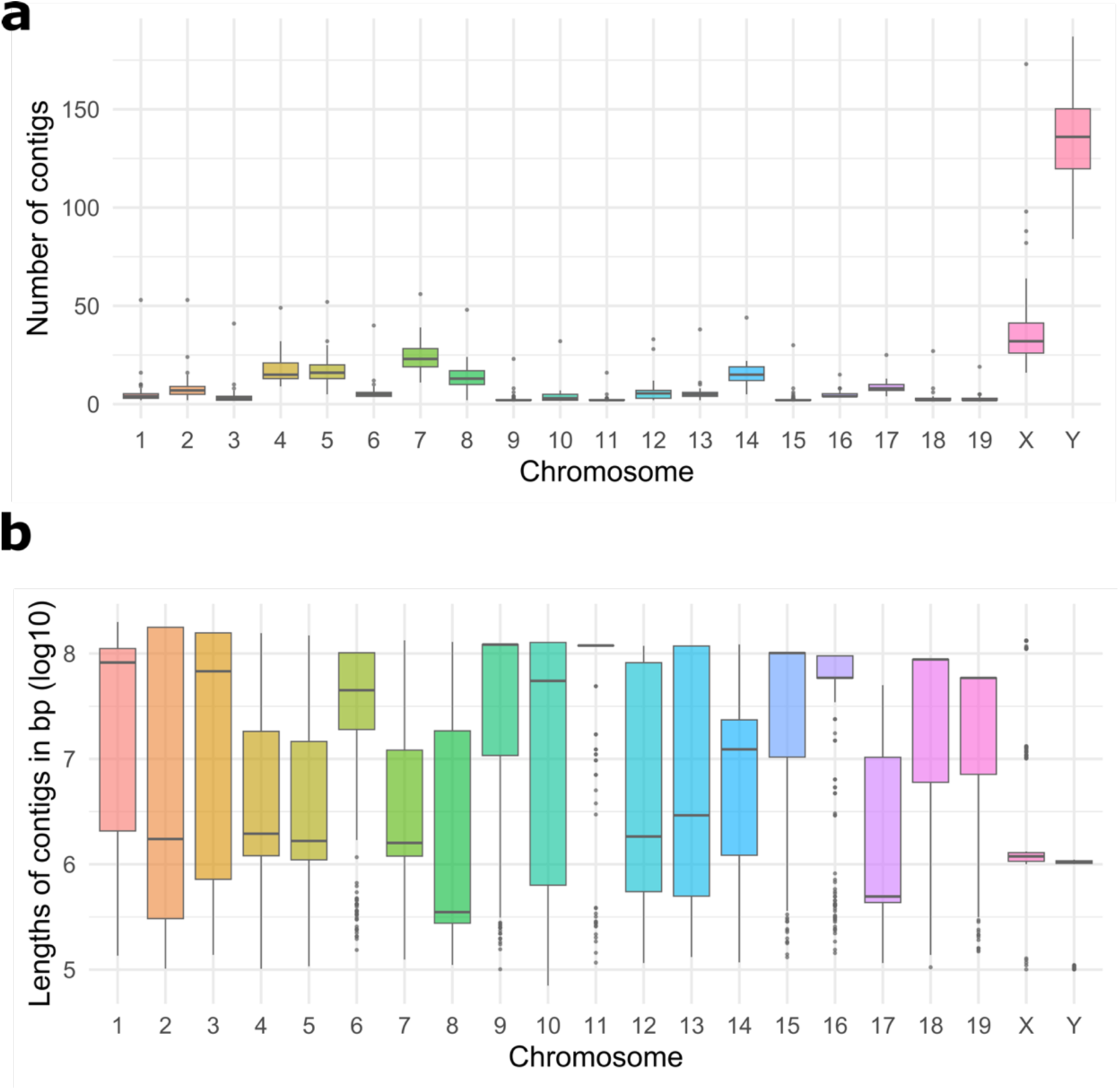
Number (a) and length (b) of contigs per chromosome. Data are pooled across all samples, excluding two low-quality samples with diploid genome map N50 values below recommended thresholds.

### Structural Variant Filtering

To assess the reproducibility and confidence of the results, in a pilot OGM run, we analyzed the degree of overlap of SVs between three technical replicates of the same biological sample. The proportion of SVs overlapping pairwise between any two replicates ranged from 80% to 88% for insertions, 60-76% for duplications, and 42-64% for deletions. Such a poor overlap of called SVs between technical replicates, especially in the case of deletions, suggests a high false positive/negative rate of detection. According to Bionano’s documentation, detected SVs of small size (below 500 bp) are generally considered unreliable. Hence, a good part of such small SVs might represent false positives. We reasoned that, if a call is found in all three technical replicates of the same biological sample, that call can be reliably considered a true event. However, if a call is detected in only one replicate sample, it is likely to be a false positive. We therefore compared the size distribution between SVs identified in all three replicates (overlapping calls) and those identified in only one replicate (non-overlapping calls; Fig. 2). Median SV size was smaller in the set of non-overlapping calls for deletions (415 bp) and insertions (428 bp) than in the set of overlapping calls (31,006 bp for deletions and 932 bp for insertions). Median size of duplications did not differ substantially between overlapping (108,518 bp) and non-overlapping (128,448 bp) calls; however, there were only 14 duplications in the non-overlapping set. These results suggest that a great deal of false positive calls can be found among insertions and deletions up to 500 bp in size. Given that the overlapping calls set also contains insertions and deletions of this size range, we re-analyzed the degree of pairwise overlap when only SVs above 500 bp are considered. The proportion of SVs overlapping between any two replicates ranged from 87% to 93% for insertions, and 67-80% for deletions. Hence, the concordance between replicates in calling insertions and deletions improves substantially when only calls above 500 bp in size are considered. We therefore excluded all calls smaller than 500 bp from all analyses of the overall effect.

**Fig. 2.**
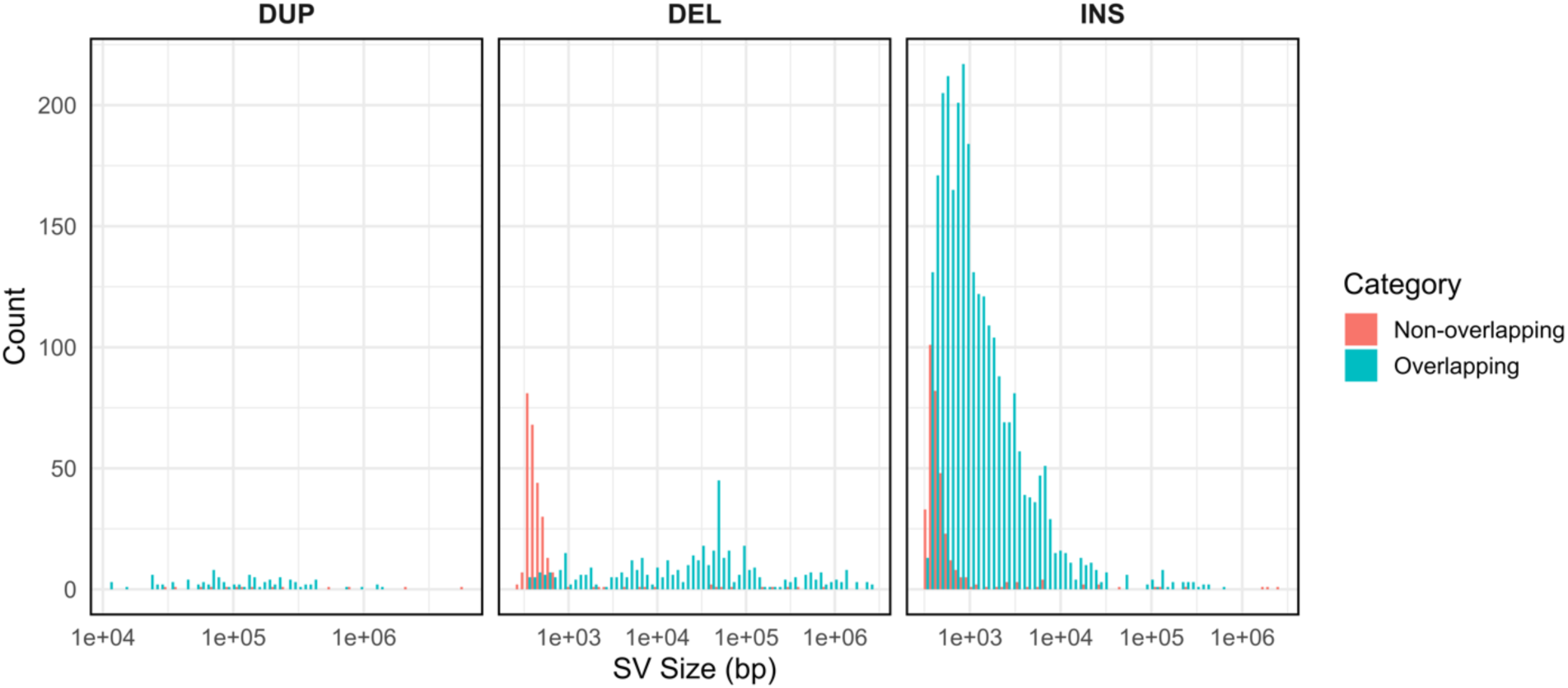
Comparison of SV sizes between variants detected in all three technical replicates (overlapping calls) and those detected in only one replicate (non-overlapping calls).

### Overall Effects

#### SV Size

Across all experiments, we observed no significant differences in SV lengths between experimental and control groups or between tissues (Supplementary Fig. S1). Because SV length distributions varied markedly across SV types (Fig. 3a), analyses were repeated separately for each type (Supplementary Fig. S2). For all SV types, within-mouse kidney-sperm differences were not significantly different from zero (paired Wilcoxon, all p > 0.06), nor did these tissue-specific differences differ between experimental and control groups (Welch’s t-test, all p > 0.19). Thus, none of the treatments measurably affected SV size.

**Fig. 3.**
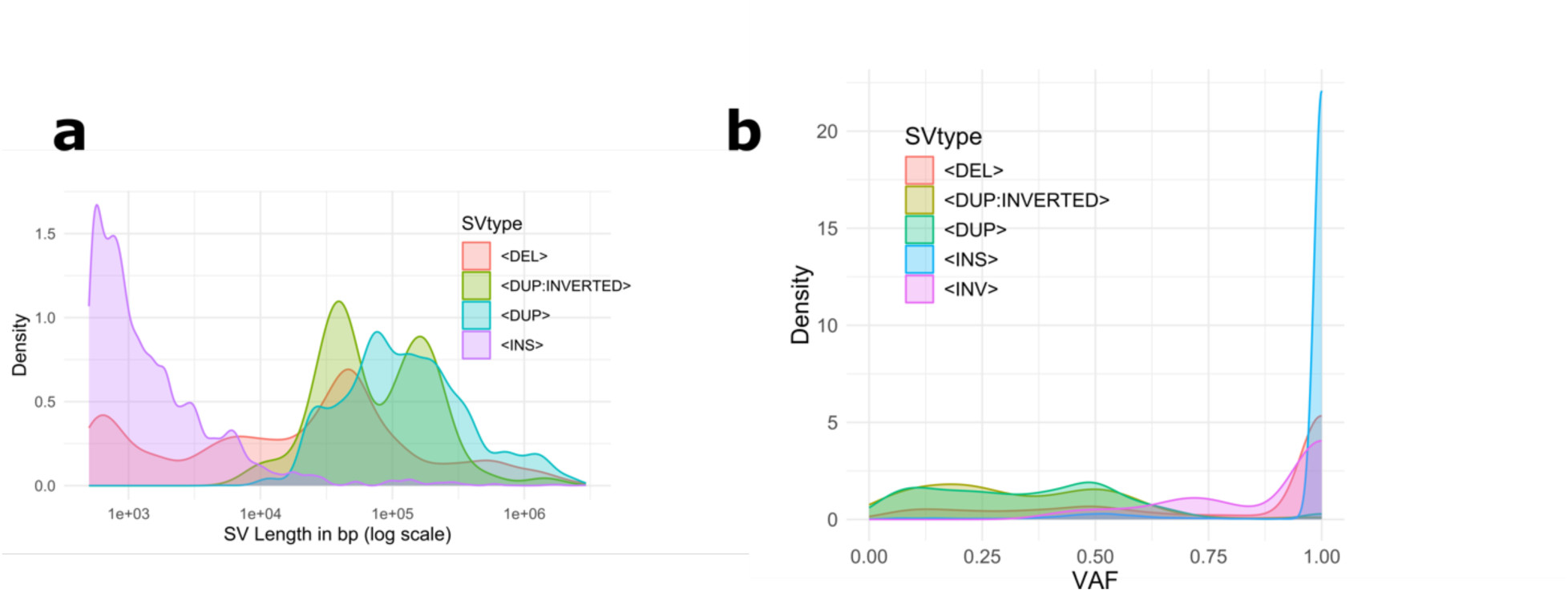
Properties of SVs by type. Distributions are shown for a) length and b) variant allele fraction.

#### SV count

We detected between 1,069 to 1,228 SVs per sample (Supplementary Fig. S3, Supplementary Table S2), among which 902 insertions, 180 deletions, 39 duplications, 2 inversions, and 2 inverted duplications, on average. The number of detected SVs did not differ significantly between experimental and control groups (Wilcoxon rank-sum, all p > 0.39). Kidney-sperm differences within mice were also non-significant (paired Wilcoxon, all p > 0.06), and the magnitude of these differences did not vary between groups (Wilcoxon rank-sum, all p > 0.22). Thus, the treatments did not alter the number of detectable SVs.

#### Variant Allele Fraction

We next compared the distribution of Variant Allele Fraction (VAF), as reported by the Bionano Solve pipeline. VAF represents the proportion of labeled DNA molecules supporting a structural variant relative to the total molecules aligned in the affected genomic region. Although VAF is not a precise quantitative measure of cellular composition, variation in VAF can provide a useful indication of subclonal mosaicism in somatic tissues or increased allelic diversity in the population of sperm cells. We found no significant differences in VAF distribution between control and experimental groups in either kidney or sperm (Supplementary Fig. S4; Wilcoxon rank-sum test, p-value > 0.08). In the experiment of ethanol consumption, statistical testing showed a borderline significant difference between the control and experimental group of sperm samples (Wilcoxon rank-sum test, p-value = 0.04); however, with a negligible effect size (rank-biserial r = 0.02). Comparison of VAF distribution between SVs of different types reveals major differences, the most pronounced between insertions and duplications - the former having high frequencies and most of the latter appearing in less than half of the molecules (Fig. 3b). We therefore examined each SV type separately. We omitted inversions and inverted duplications from this analysis due to the small number of calls (Supplementary Table S2). VAF distributions did not differ between treatment and control conditions (two-sided Wilcoxon rank-sum test, p-value > 0.05), regardless of the SV type or tissue (Supplementary Fig. S5). The only statistically significant difference was found at deletions within kidney samples in the Western diet experiment (p-value = 0.006); however, this difference was not consistent when individual mice were compared (Supplementary Fig. S6). These results suggest that the performed experiments did not affect the overall frequency of structural variation.

#### Zygosity

We next analyzed whether the experimental conditions led to shifts in the proportion of heterozygous to homozygous events. The average ratio of heterozygous to homozygous SVs in the whole dataset was 0.2 (0.15-0.28; Supplementary Table S3). We first assessed the overall distribution of homozygous to heterozygous SVs by type. We found no substantial difference in the overall distribution of SV length between homozygous and heterozygous duplications and inversions (Fig. 4). However, in the case of deletions and insertions, large events tend to be heterozygous. This is especially valid for insertions above 10^4^ bp and deletions above 10^5^ bp in size (Fig. 4). This pattern between heterozygous and homozygous events was generally similar in all experimental and control groups (Figures S7-S9), indicating no detectable treatment effects on zygosity.

**Fig. 4.**
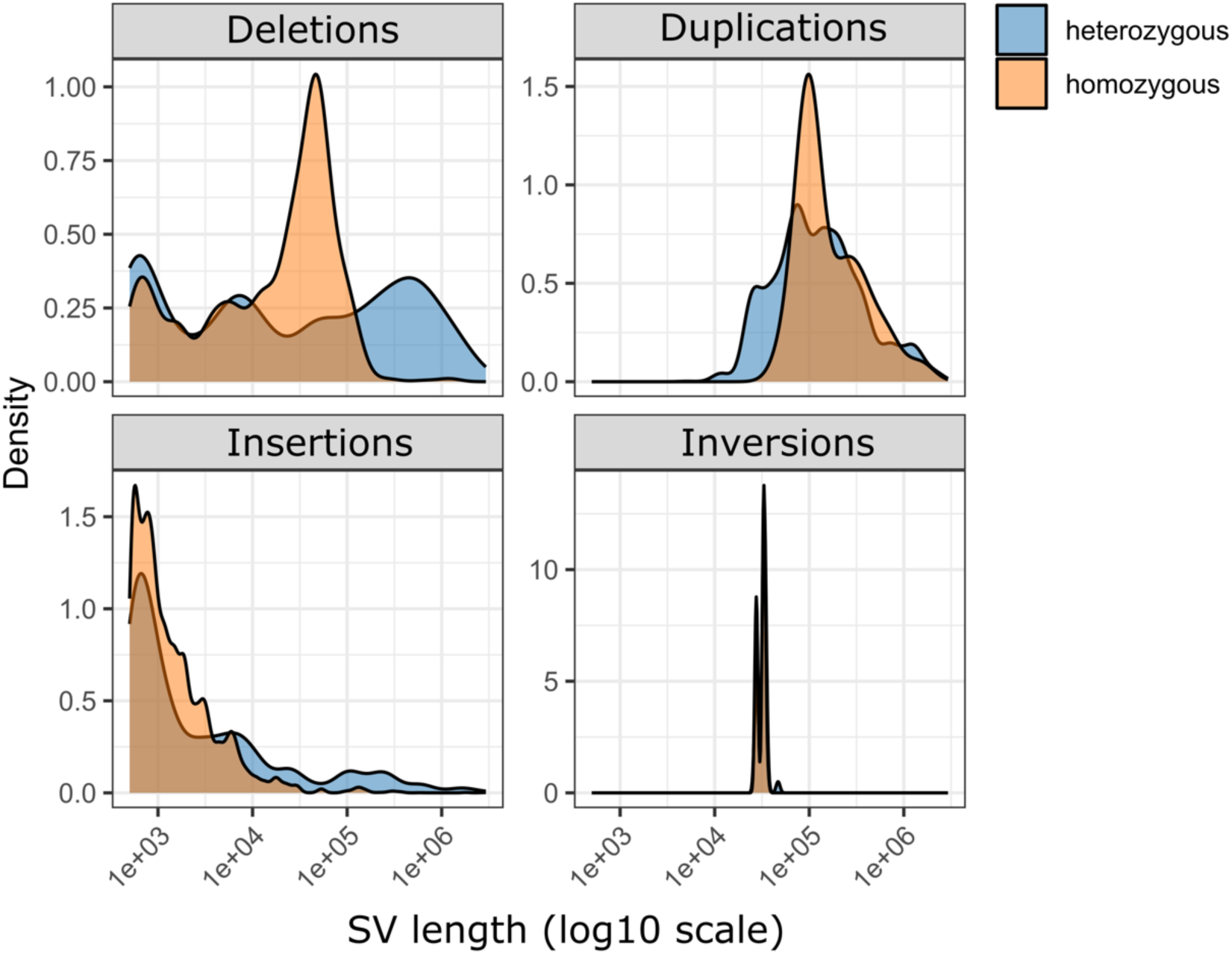
Distribution of homozygous and heterozygous variants across SV types. Inverted duplications are not shown because only two homozygous events were detected.

### De Novo Structural Variants

#### Identification and counts

To quantify de novo SV formation in the germline, we identified putative de novo events as SVs detected in sperm that did not overlap with any SVs present in the kidney sample from the same mouse. Conversely, kidney-specific de novo events were defined as SVs present in kidney but absent from sperm.

On average, 7% (5-13%) of all detected SVs were tissue-specific: 83 (52-154) per sperm and 74 (53-93) per kidney sample (Supplementary Fig. S10, Supplementary Table S4). When normalized to the calculated average VAF of 0.7, this translates to 52 de novo events per genome in the kidney and 58 in the sperm. Among the putative de novo events, the majority were insertions (51 events on average), followed by deletions (23), duplications (3), inverted duplications (1), and inversions (0.3). We observed a significant enrichment of deletions (1.8-fold, hypergeometric test, p = 1.75×10⁻^3^) and depletion of insertions (fold-change 0.80, hypergeometric test, one-sided depletion p = 5.79×10^-4^) in the de novo set relative to their proportions in the set of all detected SVs. To investigate whether the number of de novo events differed between tissues and experimental groups, we first calculated the per-mouse differences in SV counts between sperm and kidney (Δ = sperm - kidney) for each SV type (insertions, deletions, duplications, inverted duplications, inversions, and total SVs). Within each experimental group, we tested whether these within-mouse differences were significantly different from zero using paired Wilcoxon signed-rank tests. To assess whether the magnitude or direction of the sperm-kidney differences differed between experimental and control groups, we compared the Δ values between groups using Wilcoxon rank-sum tests. Across all experiments and SV types, none of the within-group or between-group comparisons reached statistical significance (Wilcoxon signed-rank test, p-values > 0.095; Wilcoxon rank-sum test, p-values > 0.15), not even when the counts were normalized to the total number of SVs per sample (Wilcoxon signed-rank test, p-values > 0.059; Wilcoxon rank-sum test, p-values > 0.21). This suggests that, in this dataset, the distribution of de novo SV counts between sperm and kidney is consistent within individual mice, and the experimental treatments did not result in detectable differences in the magnitude of sperm-kidney de novo SV count differences relative to controls.

#### Size of de novo SVs

De novo SVs were typically small, with median per-sample lengths of 560-1,401 bp (mean 5.4-104 kbp), strongly left-skewed relative to the full SV dataset. Across experiments, no differences in de novo SV length were detected between experimental and control groups in either tissue (Wilcoxon, p > 0.05). In the ethanol consumption experiment, sperm samples showed a statistically significant difference (p = 8.15×10⁻⁴), but distributions were inconsistent across individuals (Supplementary Fig. S11), indicating no reproducible treatment effect.

#### VAF and zygosity of de novo SVs

In the low-protein diet and ethanol consumption experiments, VAF distributions did not differ between groups in either tissue (Wilcoxon rank-sum test, p > 0.39). Within the Western diet experiment, we detected significant differences in the overall distribution of VAF values between the experimental (WD) and the control (C) group in sperm samples (Wilcoxon rank-sum test, p-value = 6.51 x 10^4^) but not in kidney (p-value = 0.45). However, the effect size was small (rank-biserial r = 0.121), and there was no consistency in the shape of the distribution across individual mice (Supplementary Fig. S12).

Zygosity ratios of de novo SVs (mean HET/HOM = 1.02, range 0.36-2.03; Supplementary Table S5) did not differ between groups in any experiment (Wilcoxon, all p > 0.1). Thus, the treatments did not detectably affect the allelic composition of de novo events.

### Region-Specific Effects

To identify specific genomic regions with altered SV frequency following the experiment, we focused on the Western diet experiment. For this analysis, we retained SVs of all sizes, rather than excluding events <500 bp, to avoid systematic false-negative classifications inherent to fixed-size thresholds. Variants close to such a cutoff may be retained in one sample but filtered out in another due to minor measurement differences, potentially biasing downstream statistical comparisons.

We first partitioned the genome into non-overlapping intervals based on the overlaps of SV coordinates detected in all samples from the Western diet experiment combined. For each partition, we tested whether the VAF differed significantly between the control and the experimental group. This yielded 12 partitions corresponding to ten genomic regions in which the diet potentially affected the frequency of SVs (Table 1). However, it is important to note that the statistical significance in none of these cases survives the correction for multiple testing (Benjamini-Hochberg FDR p-adjusted; Supplementary Table S6). We describe some of these regions and their association with the nearest gene. The majority of these genes are associated with obesity/overweight in human or mouse models (*PPM1A, LRMDA, ZFP503, TBX15, NEGR1, Penk, FAM155A*) or their apparent metabolic functions are relevant to high-fat diet stress (*DHRS7, Pcnx4, 6530403H02Rik*).

**Table 1.** List of candidate genomic regions.

| Partition coordinates | Median VAF |  |  |  | Genes |  | SV type |
| --- | --- | --- | --- | --- | --- | --- | --- |
|  | WD sperm | C sperm | WD kidney | C kidney | Overlapping | Nearest |  |
| chr8:9293105-9298037 | 1 | 0 | 0 | 0 | *FAM155A* | *4930453L07RIK* | INS ~400 bp |
| chr4:4179622-4185158 | 1 | 0 | 0 | 0 | - | *PENK* | INS ~400 bp |
| chr3:99528102-99542812 | 1 | 0 | 0.49 | 0 | - | *TBX15* | INS ~650 bp |
| chr3:157244435-157246145 | 1 | 0 | 0 | 0 | *NEGR1* | *4930592C13RIK* | INS ~600 bp |
| chr3:120631500-120644842 | 1 | 0 | 0 | 0 | *6530403H02RIK* | *RWDD3* | INS ~700 bp |
| chr2:3965988-3966813 | 1 | 0 | 0 | 1 | - | *FRMD4A* | INS ~15 kbp |
| chr14:6020219-6079408 | 0.11 | 0 | 0 | 0 | - | *GM3383* | DEL 398-1,206 kbp |
| chr14:6079408-6085947 | 0.14 | 0 | 0 | 0 | - | *GM3383* | DEL 398-1,206 kbp |
| chr14:6085947-6139067 | 0.14 | 0 | 0 | 0.06 | - | *GM3383* | DEL 398-1,206 kbp |
| chr14:43037134-43054481 | 0.13 | 0 | 0 | 0 | *GM6536* | *GM10376* | DUP & DEL 151-603 kbp |
| chr14:22007513-22014068 | 1 | 0 | 0 | 0 | - | *ZFP503, LRMDA* | INS ~500 bp |
| chr12:72653707-72655210 | 1 | 0 | 0 | 1 | *DHRS7* | *PCNX4, PPM1A* | INS ~500 bp |

A small insertion, approximately 600 bp in size, is detected at chr3:157244435-157246145 in the 6^th^ intron of *NEGR1* (Neuronal growth regulator 1). *NEGR1* produces a protein involved in cell adhesion, brain development, and energy balance, with variants affecting food intake, lipid storage, and metabolism. It has a role in intracellular cholesterol homeostasis, and specific SNP and CNV in *NEGR1* are linked to obesity and related traits [43–45]. Although this insertion is detected in all sperm WD samples at VAF=1, it is also detected in one C sperm and two WD kidney samples (Fig. 5).

**Fig. 5.**
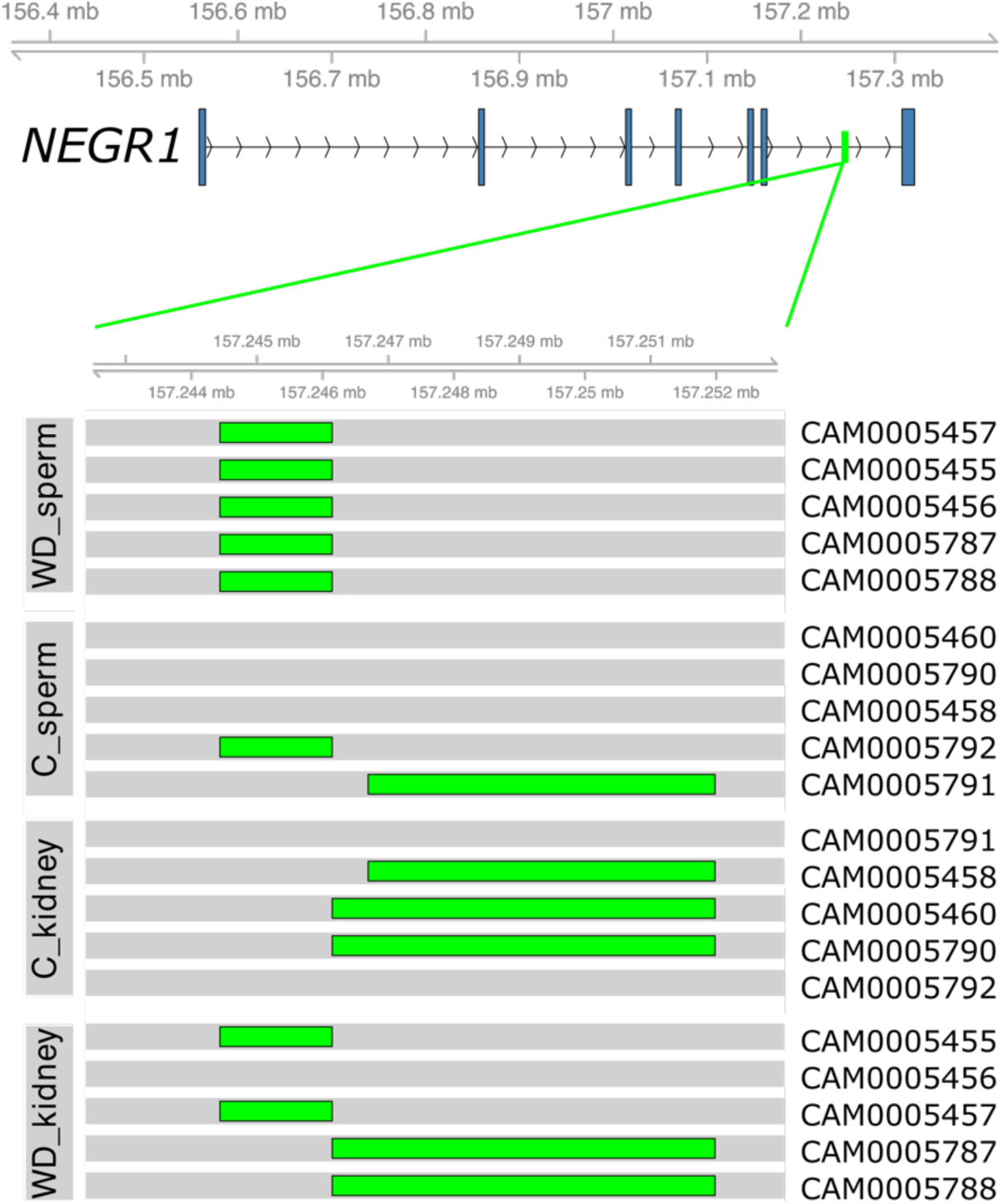
Schematic representation of the genomic region at chr3:157244435-157246145 encompassing the *NEGR1* gene. The gene structure is shown at the top, with the highlighted region enlarged below (green). The location of the insertion is indicated by a green rectangle in each sample track where it is detected. Mouse identification numbers are shown on the right, and sample groups are indicated on the left.

Another small insertion, 420-558 bp in size, is present in all sperm samples of the WD group in the region on chr14:22007496-22014725, whereas it is absent from all other samples except one kidney sample from the WD group (Fig. 6). This region is upstream of the *LRMDA* gene (Leucine Rich Melanocyte Differentiation Associated) and *ZFP503* gene (zinc finger protein 503). While *LRMDA* is primarily known for its role in pigmentation, a SNP variant near this gene was recently associated with mild obesity-related diabetes [46]. *ZFP503* (also known as *ZFHX3*) is linked to body weight regulation, with a specific variant associated with lower BMI, reduced food intake, and better metabolic health [47, 48].

**Fig. 6.**
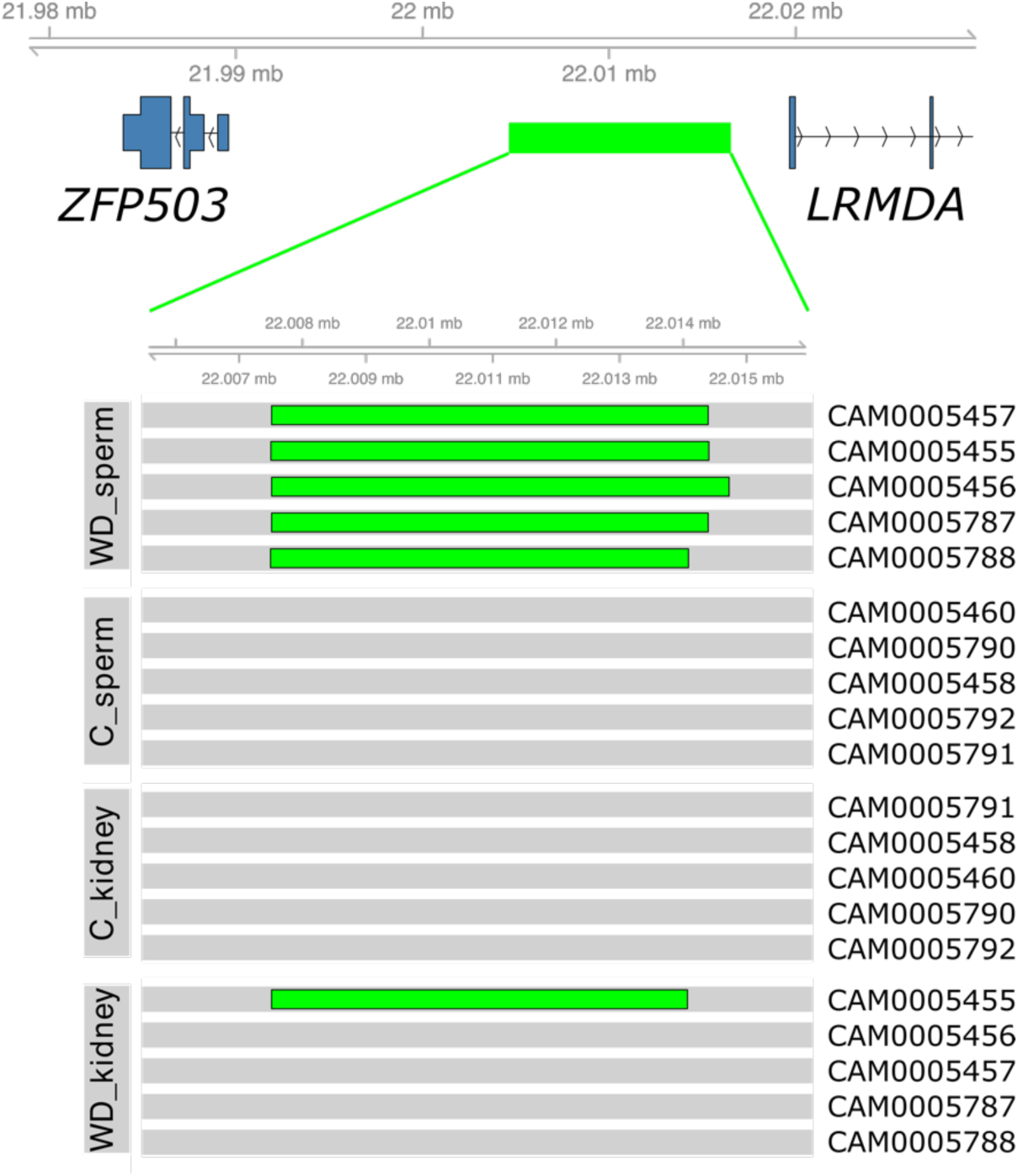
Schematic representation of the genomic region at chr14:22007496-22014725 encompassing *ZFP503* and part of the *LRMDA* gene. Gene structures are shown at the top, with the intergenic region highlighted and enlarged below (green). Green rectangles indicate the position of the insertion in sample tracks where it is detected. Mouse identification numbers are shown on the right, and sample groups are indicated on the left.

An insertion of 623-922 bp in size was detected in four WD sperm samples, one C sperm, and one WD kidney sample in the region on chr3:120631060-120646204 (Fig. 7). This region encompasses exon 10 and parts of introns 9 and 10 of the *6530403H02Rik* gene. The *6530403H02Rik* gene is classified as a long non-coding RNA (lncRNA) gene, whose downregulation has been detected in certain cell clusters of obese adipose tissue in the epididymis of mice [49].

**Fig. 7.**
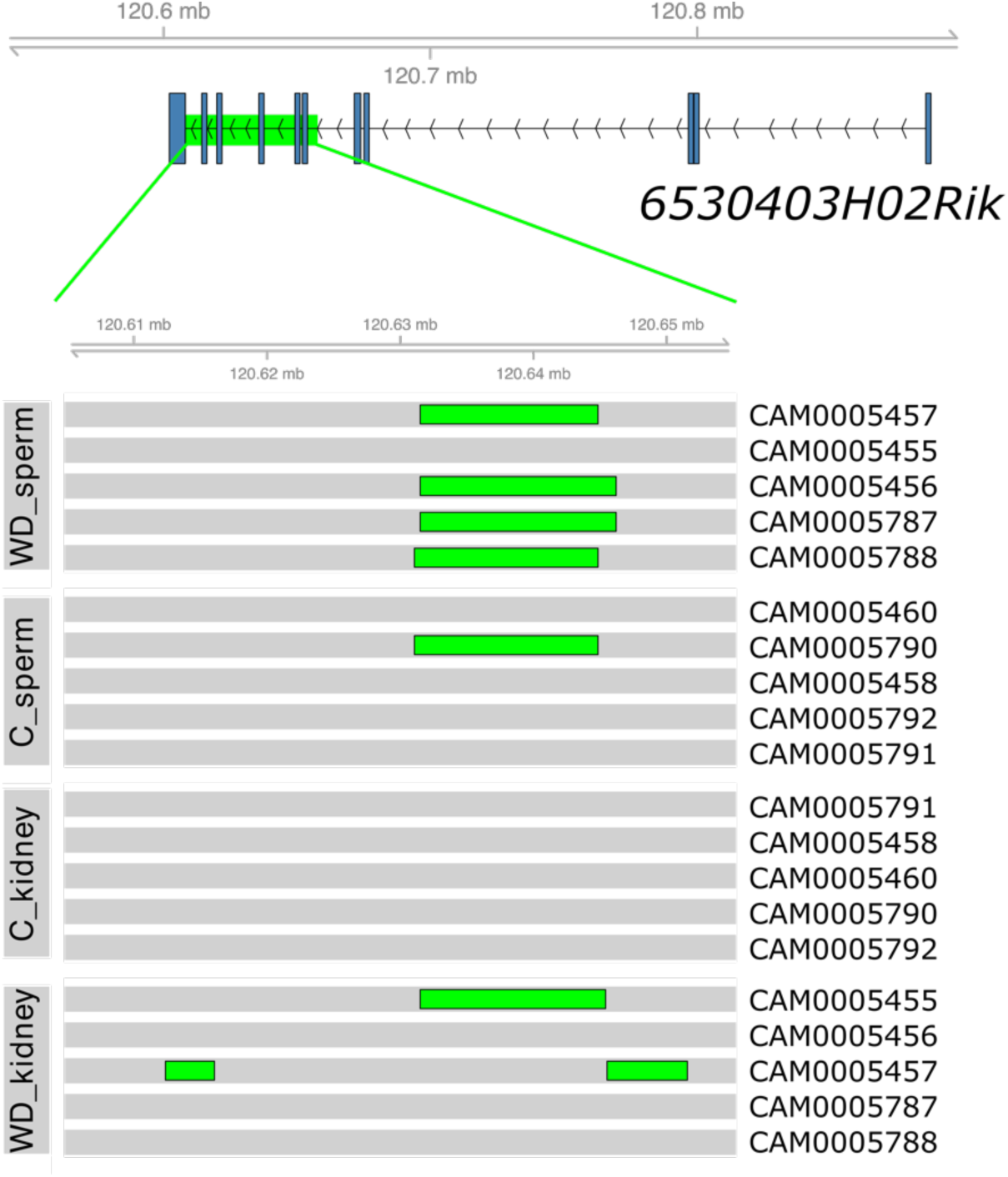
Schematic representation of the genomic region at chr3:120631060-120646204 containing the *6530403H02Rik* gene. Gene structure is shown at the top, with the highlighted region enlarged below (green). Green rectangles indicate the position of the insertion in sample tracks where it is detected. Mouse identification numbers are shown on the right, and sample groups are indicated on the left.

Another small insertion < 500 bp in size is detected in all WD sperm samples at chr8:9293105-9298037, in the first intron of *FAM155A* (Fig. 8). The gene *FAM155A* (Family with sequence similarity 155, member A) is linked to obesity, showing up in genetic studies (GWAS) as near regions associated with Body Mass Index (BMI), visceral fat, and weight gain, particularly in cancer survivors and animal models, suggesting genetic variations near *FAM155A* can influence fat distribution and obesity risk [50–52]. The same insertion is also detected in one sample from WD kidney and C kidney.

**Fig. 8.**
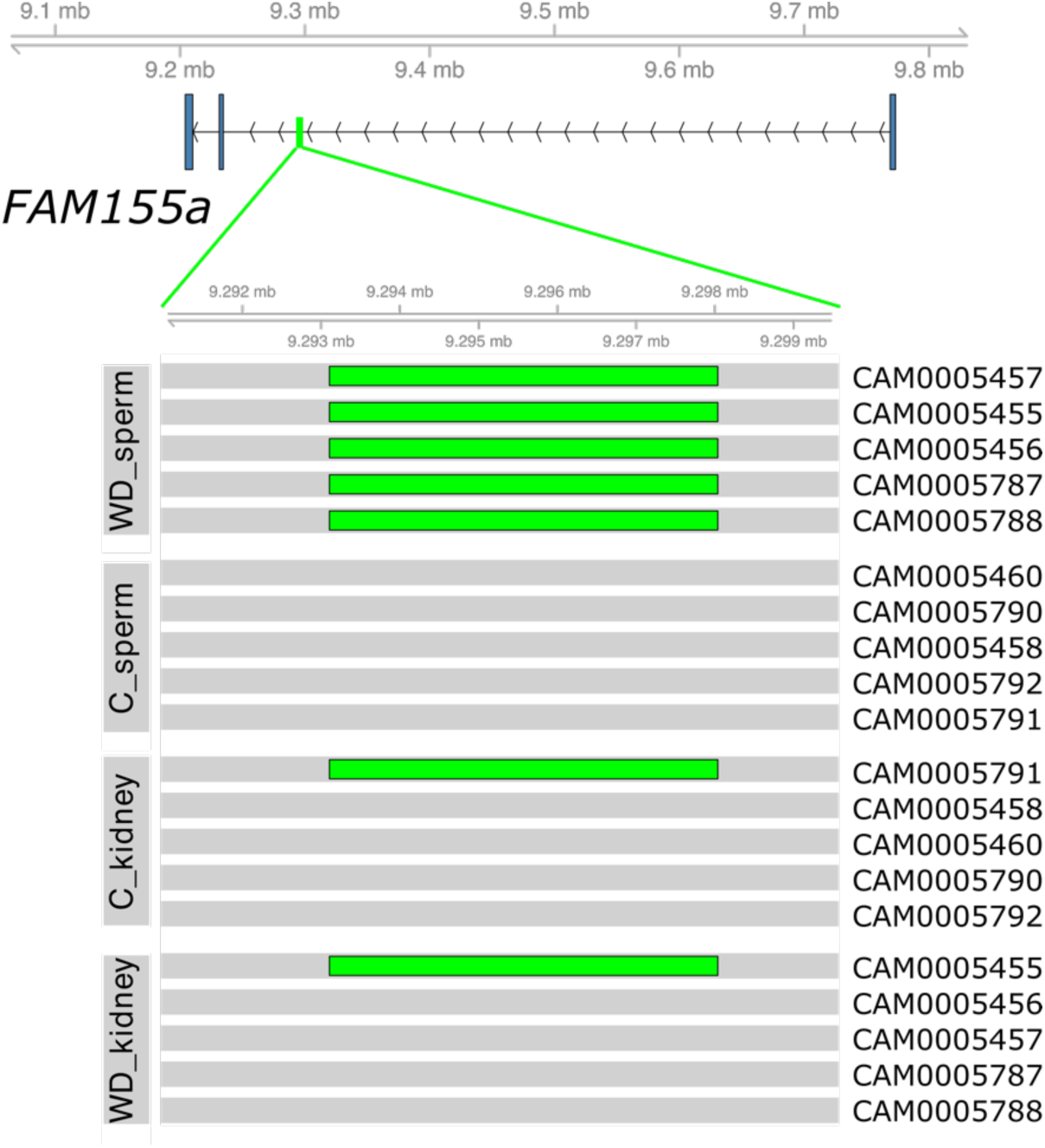
Schematic representation of the genomic region at chr8:9293105-9298037 containing the *FAM155A* gene. Gene structure is shown at the top, with the highlighted region enlarged below (green). Green rectangles indicate the position of the insertion in sample tracks where it is detected. Mouse identification numbers are shown on the right, and sample groups are indicated on the left.

## Discussion

In this study, we used optical genome mapping to investigate whether three environmental interventions - Western diet, low-protein diet, and intermittent ethanol consumption - alter structural variation (SV) in somatic tissue and the male germline. Across all experiments, we observed no evidence for widespread or systematic changes in SV landscapes, indicating that mid- to large-scale SVs remain stable under these dietary and metabolic conditions. Notably, the similar frequency of tissue-specific SVs across all groups suggests that the rate of SV formation in the germline is not elevated by these interventions. These findings stand in contrast to experimental evidence from fruit fly and yeast, where dietary composition and nutritional stress have been shown to induce CNVs and chromosomal rearrangements [18, 19]. Rather than contradicting those studies, our results suggest that environmentally induced structural plasticity may be highly context-dependent, potentially requiring specific genome architectures, life histories, or stress intensities that are not recapitulated under the conditions tested here. In mammals, robust genome maintenance mechanisms and developmental buffering may further constrain environmentally stimulated SV formation, particularly at the genome-wide level.

Across all samples, homozygous SVs were approximately five times more frequent than heterozygous SVs, consistent with expectations for genetic homogeneity in highly inbred mouse strains. In contrast, the subset of tissue-specific SVs - interpreted as putative de novo events - exhibited an approximately equal proportion of heterozygous and homozygous variants. This shift in zygosity is consistent with the recent origin of de novo mutations and supports the biological relevance of tissue-specific SVs, even though technical limitations of OGM likely inflate their absolute frequency.

Although our estimated rate of de novo SV formation in mice exceeds reported human germline rates by roughly two orders of magnitude [53, 54], this difference is unlikely to represent a genuine biological divergence and more likely reflects technical constraints inherent to SV detection, especially for smaller events. Consistent with this interpretation, the putative de novo SV set is enriched for smaller events, resulting in a markedly left-skewed size distribution relative to the full SV catalogue. Importantly, however, these technical limitations are expected to similarly affect all experimental groups, and no systematic differences were observed between treatments. Thus, while OGM imposes constraints on absolute estimates of de novo SV frequency, it is not expected to affect relative comparisons between experimental conditions.

Partition-based analyses in the Western diet experiment identified several genomic regions with nominal differences in SV frequency between treatment and control groups, some near genes implicated in metabolic regulation and associated with overweight/obesity in previous studies. Although this suggests possible local susceptibility to structural changes, none of the signals remained significant after correction for multiple testing, and most involved small SVs are close to the detection threshold. While these findings do not provide conclusive evidence for diet-induced SV changes, they highlight genomic regions that merit follow-up with higher-resolution methods and larger cohorts.

Together, these results provide an important constraint on models proposing that common environmental exposures may broadly elevate (germline) structural mutation rates [26, 38, 55]. While environmentally induced SV formation is clearly possible in some systems and may occur at specific vulnerable loci [18, 26, 56], our findings indicate that such effects, if present in mammals, are subtle, highly localized, or below the detection limits of current genome-wide approaches. Instead, SV responses to environmental exposures more likely operate on population scales through natural selection, by drawing from standing genetic variation as suggested by population-scale evolutionary genomics studies [39, 40, 57, 58].

By explicitly reporting the absence of strong effects, this study contributes to a more balanced understanding of environmental influences on genome stability. It suggests that environmentally induced phenotypic and transgenerational effects observed in mammals are more likely mediated through epigenetic regulation, chromatin dynamics, or small RNA pathways rather than through large-scale changes in genome structure. Future studies combining larger sample sizes, higher-resolution SV detection, and multigenerational designs will be essential for determining whether environmentally induced structural variation plays a measurable role in mammalian evolution or disease risk.

## Methods

### Ethics approval

All procedures involving mice were approved by the Ministry of Agriculture, Forestry and Water Management of the Republic of Croatia, Class: UP/I-322-01/19-01/59 and case number: 525-10/0543-20-4, and carried out in compliance with relevant guidelines: the ILAR Guide for the Care and Use of Laboratory Animals, the EU Directive 2010/63/EU, the Croatian Animal Welfare Act (NN 102/17), and the Ruđer Bošković Institute’s policy on animal use and ethics. Mice were humanely euthanized by cervical dislocation. The authors complied with the ARRIVE guidelines.

### Experimental Design

Prior to the start of each experiment, mice were maintained under standard laboratory conditions at the Ruđer Bošković Institute (Zagreb, Croatia). Food and water were provided ad libitum for all animals before and during the experiments.

For the low-protein diet experiment, male C57BL/6 mice aged 4-5 weeks were randomly assigned to the control (C) or experimental group (LPD). Five mice in the LPD group were fed a diet with reduced protein content, containing 8.8% proteins, 8.1% fat, and 11% sucrose (Low Protein diet, catalog #E15202-24; ssniff Spezialdiäten GmbH, Germany), and three control mice were fed standard laboratory chow (4RF 21; Mucedola srl, Italy). The experiment lasted 80 days.

For the ethanol consumption experiment, male C57BL/6 mice aged 8-9 weeks were singly housed and maintained on a reverse 12 h light / 12 h dark cycle. Five mice in the experimental (EtOH) group received intermittent access to 15% (v/v) ethanol for 2 h at the onset of each dark phase, according to the free-choice drinking model (Griffin 2014). Four mice from the control group (C) were kept under the same conditions without access to alcohol. Mice from both groups were fed standard laboratory chow. The experiment lasted 40 days.

### Tissue Collection, DNA Preparation, and Optical Genome Mapping

Tissue collection and high-molecular-weight (HMW) DNA extractions were performed from mice in the low-protein diet and ethanol consumption experiments as described in [42]. Optical genome mapping was performed on the Saphyr platform (Bionano Genomics) at the Institute of Applied Biotechnologies a.s. (IAB, Olomouc, Czech Republic) following the procedures described in [42]. For each mouse, OGM was performed on HMW DNA isolated from sperm and kidney tissue.

In this study, we also used OGM data generated from mice in the previously conducted Western diet experiment [42] (PRJCA043930, Genome Sequence Archive). We generated optical genome maps from sperm and kidney of one additional mouse in the control group of this experiment.

### Data Filtering

De novo assembly was performed using Bionano Solve 3.8.1. Summary statistics from Informatics Report files (exp_informaticsReportSimple.txt) were used to assess data and assembly quality. The number and size of contigs per chromosome were obtained from the file exp_refineFinal1_merged.xmap within the output/contigs/annotation/ directory.

### Parsing VCF Files

SVs were extracted from variants_combine_filters_inMoleRefine1.ogm.vcf files generated by Bionano Solve. Only SVs identified by the SMAP (SV) caller were retained; CNV-caller-derived events were excluded due to low breakpoint precision and lack of variant allele frequency (VAF) information. Translocation calls were excluded because the VCF reports only one-sided breakpoints, often without a reliable mate coordinate. For insertions and deletions, breakpoint uncertainty was addressed by expanding the annotated coordinates using CIPOS and CIEND values, representing the maximum confidence-interval boundaries. This adjustment was not necessary for duplications, inverted duplications, or inversions, which have precisely defined breakpoints. All SVs were converted into BED format for downstream analyses.

### Pilot Run

To assess the reproducibility of SV detection, a single sperm-derived HMW DNA sample was divided into three aliquots, independently labeled with DLE-1, processed by optical mapping, and analyzed using the filtering steps described above. Following the above-described procedure for removal of CNVs, sex-chromosome SVs, and translocations, and the correction of VCF coordinates, call datasets were divided based on their classification into three major SV types: deletions, duplications, and insertions. Because the total numbers of inversions and inverted duplications were very low (7 and 3 in the total dataset, respectively), these classes were excluded from analyses. For each of the three SV type datasets, bedtools intersect [59] with the 50% reciprocal overlap criterion, and the *-u* flag was used to find the overlapping SVs in a pairwise manner. The proportion of overlapping events for each replicate A with another replicate B was determined as the number of overlapping events in A divided by the total number of events detected in A. For comparison of SV size distributions, SVs were further separated into overlapping calls (any overlap between all three replicate samples) and non-overlapping calls (having no overlap in the other two replicate samples).

### Comparisons of SV Sizes

SV lengths were extracted from the SVLEN field in the VCF file. Because SVLEN is not reported for inversions, inversion sizes were calculated as the absolute genomic distance between REFPOS and REFEND coordinates. For each mouse and each SV type, the median SV length was computed separately for sperm and kidney. Within each mouse, the kidney-sperm difference in median SV length was then calculated. Paired Wilcoxon tests were used to compare differences within groups, and unpaired Welch t-tests were used to compare kidney-sperm differences between experimental and control groups.

### Zygosity Assignment

Zygosity is annotated by the Bionano Solve pipeline in the ZYG field of the VCF file for deletions, insertions, and inversions, but not for duplications or inverted duplications (“unknown”). To standardize analyses across SV classes, we classified all SVs with VAF < 0.97 as heterozygous and all SVs with VAF ≥ 0.97 as putative homozygous.

### Genomic Partition

Using all SVs from the Western diet experiment, the genome was partitioned into 10,791 non-overlapping intervals using *bedops* [60]. These partitioned intervals were then intersected with individual samples’ SV coordinates to assign VAF value from each sample to every partition, by using *bedtools intersect −loj*. In cases where a single partition overlapped more than one SV within the same genome (sample), the average VAF value of all overlapping SVs was reported for that partition in that sample, by using *bedtools groupby*. Intervals without SVs were assigned a VAF of 0. For each genomic partition >500 bp, VAF values were compared between WD and control samples in sperm using a two-sided Mann-Whitney U test (Wilcoxon rank-sum test). Median VAFs were calculated for each group (WD_sperm, C_sperm, WD_kidney, and C_kidney) and reported for descriptive comparison. Only partitions with statistically significant difference (p-value < 0.05) were considered. The difference between sperm and kidney in the median VAF was calculated for each partition, and only those with median VAF in sperm higher than in kidney were retained.

## Acknowledgements

Data analysis was performed on the high-performance computing cluster at the University Computing Centre (SRCE), University of Zagreb. We thank Ranko Stojković for mouse facility support, Petra Mikolčević for help with preparing sperm cells, and Regina Bezděková Fillerová for help with optical genome mapping.

## Author contributions

I.P.: Data curation, Investigation, Methodology, Writing - review and editing. Ž.P.: Conceptualization, Funding acquisition, Investigation, Methodology, Formal analysis, Project administration, Supervision, Visualization, Writing - original draft, Writing - review and editing. All authors have read and approved the final manuscript.

## Additional information

### Consent for publication

Not applicable.

## References

1. Chaisson, M. J. P. et al. Resolving the complexity of the human genome using single-molecule sequencing. Nature. 517, 608–11 (2015).

2. Sudmant, P.H. et al. An integrated map of structural variation in 2,504 human genomes. Nature. 526, 75–81 (2015).

3. Gonzalez, E. et al. The influence of CCL3L1 gene-containing segmental duplications on HIV-1/AIDS susceptibility. Science. 307, 1434–1440 (2005).

4. Perry, G. H. et al. Diet and the evolution of human amylase gene copy number variation. Nat. Genet. 39, 1256–1260 (2007).

5. Pezer, Ž., Harr, B., Teschke, M., Babiker, H. & Tautz, D. Divergence patterns of genic copy number variation in natural populations of the house mouse (*Mus musculus domesticus*) reveal three conserved genes with major population-specific expansions. Genome Res. 25, 1114–24 (2015).

6. Carvalho, C. M. & Lupski, J. R. Mechanisms underlying structural variant formation in genomic disorders. Nat. Rev. Genet. 17, 224–238 (2016).

7. Dorant, Y. et al. Copy number variants outperform SNPs to reveal genotype-temperature association in a marine species. Mol. Ecol. 29, 4765–4782 (2020).

8. Pokrovac, I. & Pezer, Ž. Recent advances and current challenges in population genomics of structural variation in animals and plants. Front. Genet. 13, 1060898; doi: 10.3389/fgene.2022.1060898. (2022).

9. Zhang, F., Gu, W., Hurles, M. E. & Lupski, J. R. Copy number variation in human health, disease, and evolution. Annu. Rev. Genomics. Hum. Genet. 10, 451–481 (2009).

10. Lee, C. & Scherer, S. W. The clinical context of copy number variation in the human genome. Expert. Rev. Mol. Med. 12, e8; doi: 10.1017/S1462399410001390. (2010).

11. Thomas, N. S., et al. De novo apparently balanced translocations in man are predominantly paternal in origin and associated with a significant increase in paternal age. J. Med. Genet. 47, 112–5 (2010).

12. Somers, C. M., Yauk, C. L., White, P. A., Parfett, C. L. & Quinn, J. S. Air pollution induces heritable DNA mutations. Proc. Natl. Acad. Sci. 99, 15904–15907 (2002).

13. Rogstad, S. H., Keane, B. & Collier, M. H. Minisatellite DNA mutation rate in dandelions increases with leaf-tissue concentrations of Cr, Fe, Mn, and Ni. Environ. Toxicol. Chem. 22, 2093–2099 (2003).

14. Marchetti, F. et al. Sidestream tobacco smoke is a male germ cell mutagen. Proc. Natl. Acad. Sci. 108, 12811–12814 (2011).

15. Adewoye, A. B., Lindsay, S. J., Dubrova, Y. E. & Hurles, M. E. The genome-wide effects of ionizing radiation on mutation induction in the mammalian germline. Nat. Commun. 6, 6684; doi: 10.1038/ncomms7684. (2015).

16. Chain, F. J. J., Flynn, J. M., Bull, J. K. & Cristescu, M. E. Accelerated rates of large-scale mutations in the presence of copper and nickel. Genome Res. 29, 64–73 (2019).

17. Arlt, M. F., Wilson, T. E. & Glover, T. W. Replication stress and mechanisms of CNV formation. Curr. Opin. Genet. Dev. 22, 204–210 (2012).

18. Aldrich, J. C. & Maggert, K. A. Transgenerational inheritance of diet-induced genome rearrangements in Drosophila. PLoS Genet. 11, e1005148; doi: 10.1371/journal.pgen.1005148. (2015).

19. Kroll, E. et al. Starvation-associated genome restructuring can lead to reproductive isolation in yeast. PLoS One. 8, e66414; doi: 10.1371/journal.pone.0066414. (2013).

20. Zhang, F., Carvalho, C. M. & Lupski, J. R. Complex human chromosomal and genomic rearrangements. Trends. Genet. 25, 298–307 (2009).

21. Hastings, P. J., Lupski, J. R., Rosenberg, S. M. & Ira, G. Mechanisms of change in gene copy number. Nat. Rev. Genet. 10, 551–564 (2009).

22. Aguilera, A. & Garcia-Muse, T. Causes of genome instability. Annu. Rev. Genet. 47, 1–32 (2013).

23. Gaillard, H. & Aguilera, A. Transcription as a threat to genome integrity. Annu. Rev. Biochem. 85, 291–317 (2016).

24. Wilson, T. E. et al. Large transcription units unify copy number variants and common fragile sites arising under replication stress. Genome Res. 25, 189–200 (2015).

25. Saponaro, M. et al. RECQL5 controls transcript elongation and suppresses genome instability associated with transcription stress. Cell. 157, 1037–1049 (2014).

26. Hull, R. M., Cruz, C., Jack, C. V. & Houseley, J. Environmental change drives accelerated adaptation through stimulated copy number variation. PLoS Biology. 15, e2001333; doi: 10.1371/journal.pbio.2001333. (2017).

27. Rando, O. J. & Verstrepen, K. J. Timescales of genetic and epigenetic inheritance. Cell. 128, 655–668 (2007).

28. Hamperl, S. & Cimprich, K. A. A. Conflict Resolution in the Genome: How Transcription and Replication Make It Work. Cell. 167, 1455–1467 (2016).

29. Palmer, N. O., Bakos, H. W., Owens, J., Setchell, B. P. & Lane, M. Diet and exercise in an obese mouse fed a high fat diet improves metabolic health and reverses perturbed sperm function. Am. J. Physiol. Endocrinol. Metab. 5005, 768–780 (2012).

30. Donkin, I. & Barrès, R. Sperm epigenetics and influence of environmental factors. Mol. Metab. 14, 1–11 (2018).

31. Carone, B. R. et al. Paternally induced transgenerational environmental reprogramming of metabolic gene expression in mammals. Cell. 143, 1084–1096 (2010).

32. Anderson, L. M. et al. Preconceptional fasting of fathers alters serum glucose in offspring of mice. Nutrition. 22, 327–331 (2006).

33. de Castro Barbosa, T., et al. High-fat diet reprograms the epigenome of rat spermatozoa and transgenerationally affects metabolism of the offspring. Mol. Metab. 5, 184–197 (2015).

34. Chen, Q. et al. Sperm tsRNAs contribute to intergenerational inheritance of an acquired metabolic disorder. Science. 351, 397–400 (2016).

35. Finegersh, A. & Homanics, G. E. Paternal alcohol exposure reduces alcohol drinking and increases behavioral sensitivity to alcohol selectively in male offspring. PLoS One. 9, e99078; doi: 10.1371/journal.pone.0099078. (2014).

36. Gapp, K. et al. Implication of sperm RNAs in transgenerational inheritance of the effects of early trauma in mice. Nat. Neurosci. 17, 667–669 (2014).

37. Liu, S., Holmes, A. D., Gupta, A., Katzman, S. & Sharma, U. Epididymal dynamics and preimplantation roles of a sperm-enriched 5’ fragment of tRNA-valine. Cell Rep. 44, 116366; doi: 10.1016/j.celrep.2025.116366. (2025).

38. Skinner, M. K., Guerrero-Bosagna, C. & Haque, M. M. Environmentally induced epigenetic transgenerational inheritance of sperm epimutations promote genetic mutations. Epigenetics. 10, 762–71 (2015).

39. Bolognini, D. et al. Recurrent evolution and selection shape structural diversity at the amylase locus. Nature. 634, 617–625 (2024).

40. Pokrovac, I., Rohner, N. & Pezer, Ž. The prevalence of copy number increase at multiallelic copy number variants associated with cave colonization. Mol. Ecol. 33, e17339.; doi: 10.1111/mec.17339. (2024).

41. Bruder, C. E. et al. Phenotypically concordant and discordant monozygotic twins display different DNA copy-number-variation profiles. Am. J. Hum. Genet. 82, 763–771 (2008).

42. Pokrovac, I., Stipoljev, S. & Pezer, Ž. TelOMpy enables single-molecule resolution of telomere length from optical genome mapping data. BMC Biol. 23, 336; doi: 10.1186/s12915-025-02437-y. (2025).

43. Kim, H. et al. The new obesity-associated protein, neuronal growth regulator 1 (NEGR1), is implicated in Niemann-Pick disease Type C (NPC2)-mediated cholesterol trafficking. Biochem. Biophys. Res. Commun. 482, 1367–1374 (2017).

44. Willer, C. J. et al. Six new loci associated with body mass index highlight a neuronal influence on body weight regulation. Nat. Genet. 41, 25–34 (2009).

45. Speliotes, E. K. et al. Association analyses of 249,796 individuals reveal 18 new loci associated with body mass index. Nat. Genet. 42, 937–48 (2010).

46. Mansour Aly, D., et al. Genome-wide association analyses highlight etiological differences underlying newly defined subtypes of diabetes. Nat. Genet. 53, 1534–1542 (2021).

47. Turcot, V. et al. Protein-altering variants associated with body mass index implicate pathways that control energy intake and expenditure in obesity. Nat. Genet. 50, 26–41 (2018).

48. Nolan, P. M. et al. A missense mutation in zinc finger homeobox-3 (ZFHX3) impedes growth and alters metabolism and hypothalamic gene expression in mice. FASEB J. 37, e23189; doi: 10.1096/fj.202201829R. (2023).

49. Lu, Z., Ding, L., Tian, X. & Wang, Q. Single cell RNA-sequencing data generated from mouse adipose tissue during the development of obesity. Data Brief. 53, 110119; doi: 10.1016/j.dib.2024.110119. (2024).

50. Kim, H. J. et al. A genome-wide association study on abdominal adiposity-related traits in adult Korean men. Obes. Facts. 15, 590–599 (2022).

51. Wilson, C. L. et al. Genetic and clinical factors associated with obesity among adult survivors of childhood cancer: A report from the St. Jude Lifetime Cohort. Cancer. 121, 2262–70 (2015).

52. Cai, Z., Christensen, O. F., Lund, M. S., Ostersen, T. & Sahana, G. Large-scale association study on daily weight gain in pigs reveals overlap of genetic factors for growth in humans. BMC Genomics. 23, 133; doi: 10.1186/s12864-022-08373-3. (2022).

53. Belyeu, J. R. et al. De novo structural mutation rates and gamete-of-origin biases revealed through genome sequencing of 2,396 families. Am. J. Hum. Genet. 108, 597–607 (2021).

54. Collins, R. L. et al. A structural variation reference for medical and population genetics. Nature. 581, 444–451 (2020).

55. Yona, A. H., Frumkin, I. & Pilpel, Y. A relay race on the evolutionary adaptation spectrum. Cell. 163, 549–559 (2015).

56. Wirbisky, S. E. & Freeman, J. L. Atrazine exposure elicits copy number alterations in the zebrafish genome. Comp. Biochem. Physiol. C. Toxicol. Pharmacol. 194, 1–8 (2017).

57. Jones, F. C. et al. The genomic basis of adaptive evolution in threespine sticklebacks. Nature. 484, 55–61 (2012).

58. Zong, S.B., Li, Y.L., & Liu, J.X. Genomic architecture of rapid parallel adaptation to fresh water in a wild fish. Mol. Biol. Evol. 38, 1317–1329 (2021).

59. Quinlan, A. R. & Hall, I. M. BEDTools: a flexible suite of utilities for comparing genomic features. Bioinformatics. 26, 841–2 (2010).

60. Neph, S. et al. BEDOPS: high-performance genomic feature operations. Bioinformatics. 28, 1919–20 (2012).

