## Supplementary Fig. for "Stability of germline structural variation under dietary and ethanol challenges"

**Table S1**. OGM quality check metrics
**Table S2**. Number of detected SVs per sample
**Table S3.** SV counts by zygosity
**Table S4.** De novo SV counts
**Table S5.** De novo SV counts by zygosity
**Table S6.** Candidate regions with differences in SV frequency
**Fig. S1.** Distribution of SV size by sample group
**Fig. S2.** Comparison of SV size distribution between sample groups and by SV type
**Fig. S3.** Number of detected SVs per mouse
**Fig. S4.** Distribution of variant allele fraction
**Fig. S5.** Comparisons of VAF distributions between groups and by SV type
**Fig. S6.** Comparison of VAF distribution of deletions between kidney samples of individual mice in the Western Diet experiment
**Fig. S7.** Distribution of homozygous and heterozygous SVs in the Western diet experiment
**Fig. S8.** Distribution of homozygous and heterozygous SVs in the low-protein diet experiment
**Fig. S9.** Distribution of homozygous and heterozygous SVs in the ethanol consumption experiment
**Fig. S10.** Number of detected de novo SVs per mouse
**Fig. S11.** Distribution of lengths of tissue-specific de novo SVs
**Fig. S12.** Distribution of variant allele fraction of tissue-specific de novo SVs

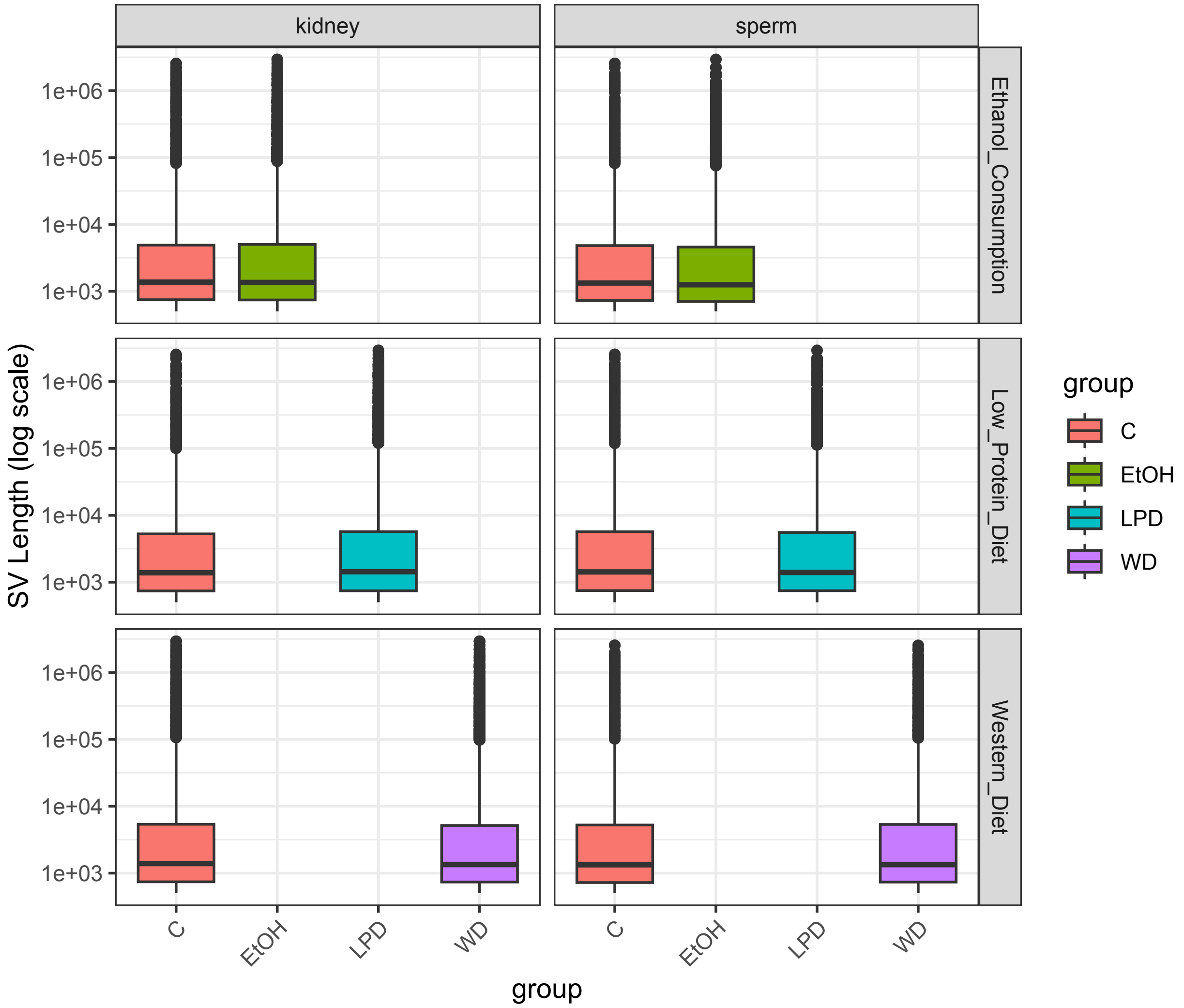


**Fig. S1.** Distribution of SV sizes by sample group.

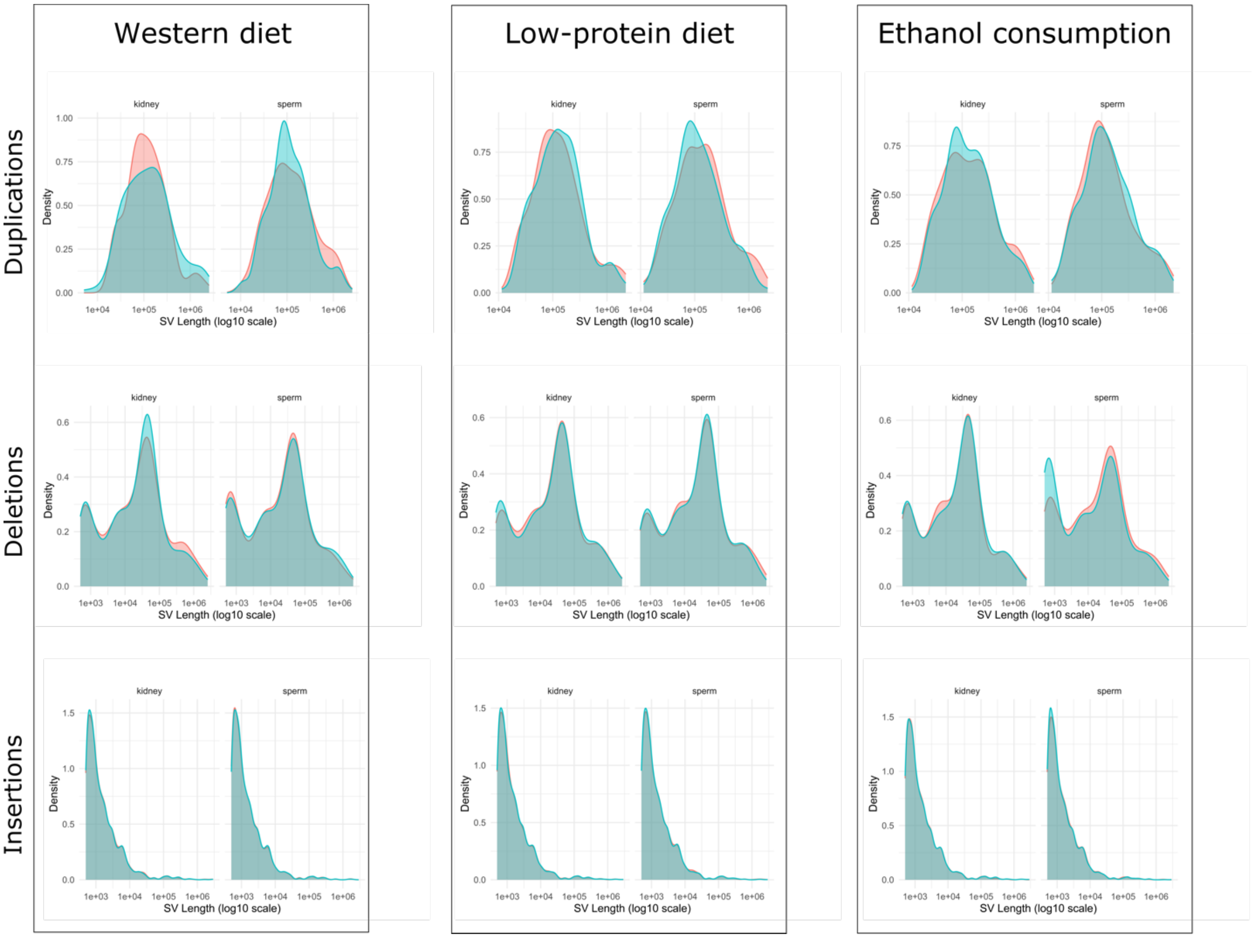


**Fig. S2.** Comparison of SV size distribution between sample groups and by SV type. Data from the control groups is shown in red and from the experimental groups in green.

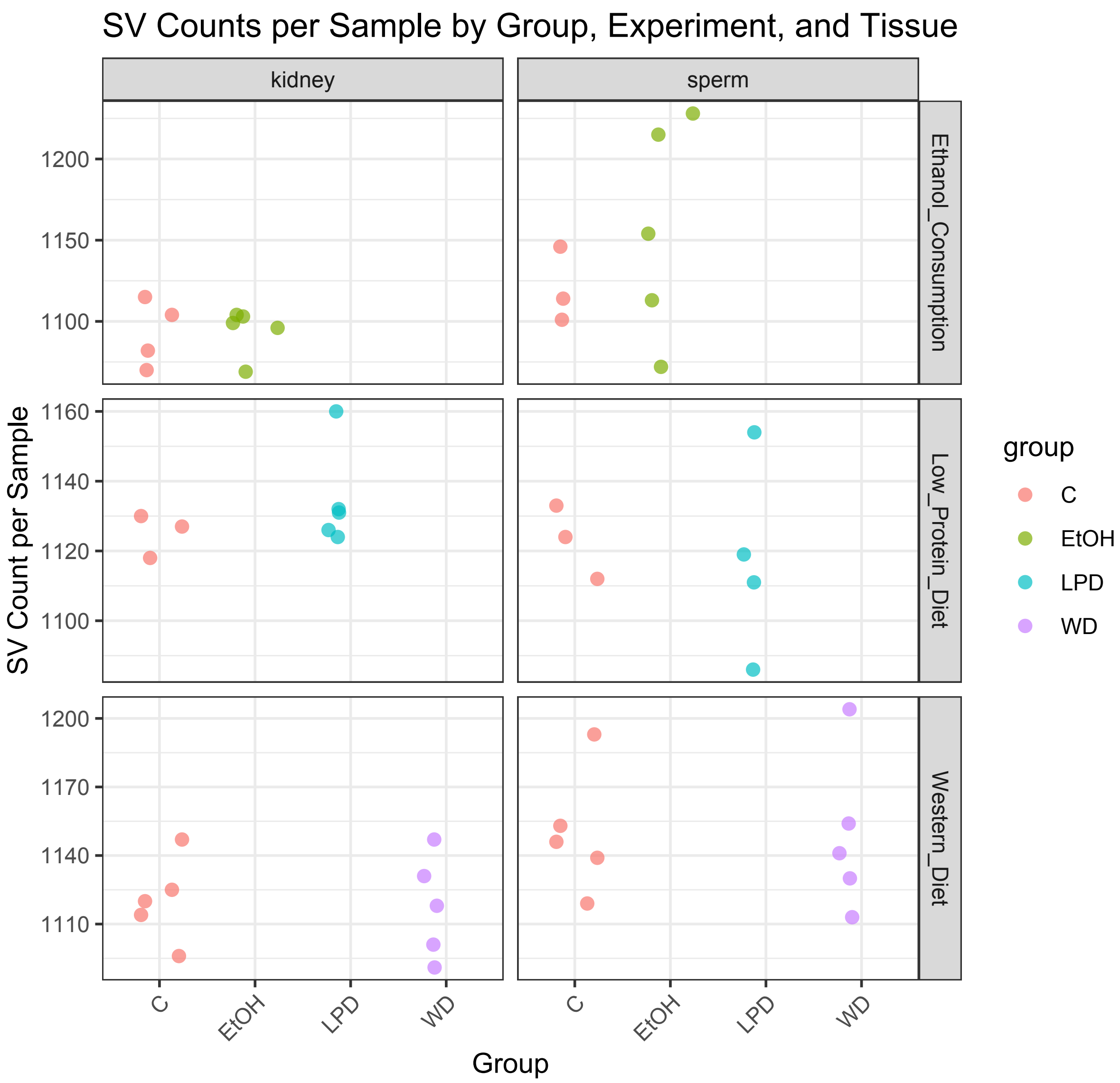


**Fig. S3.** Number of detected SVs per mouse

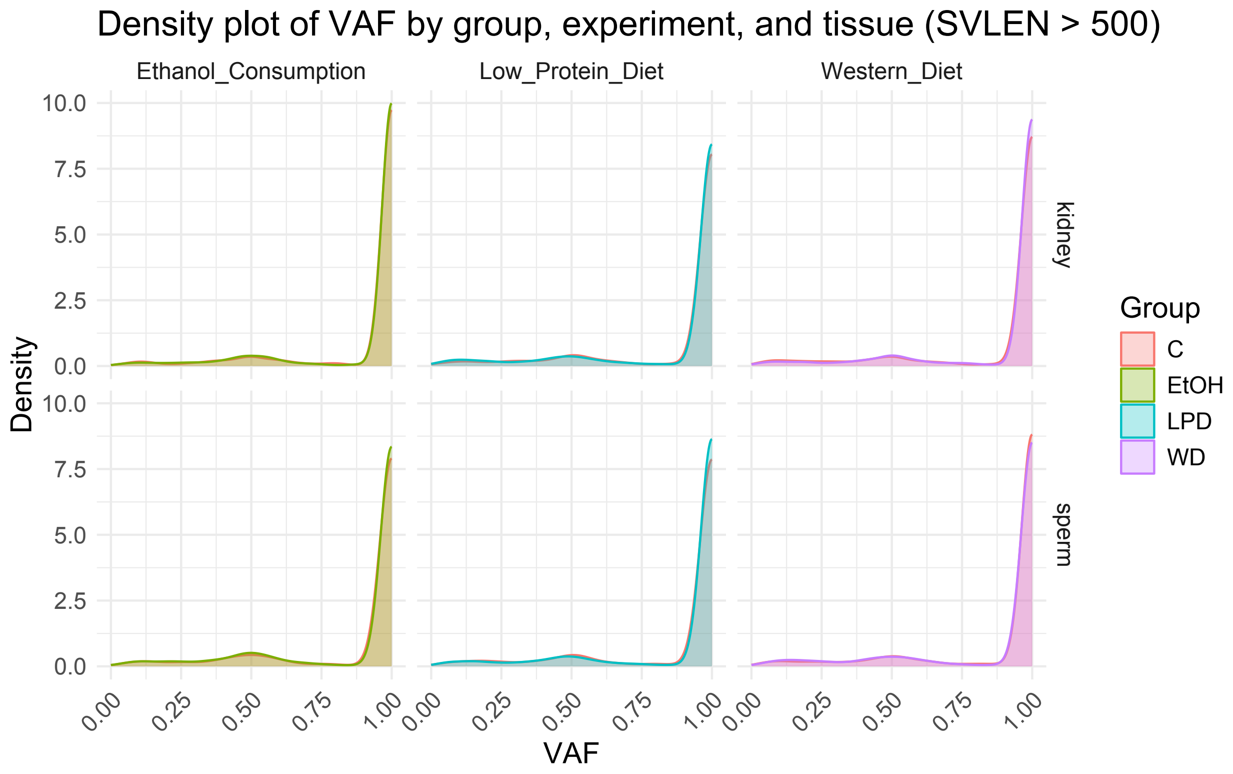


**Fig. S4.** Distribution of variant allele fraction

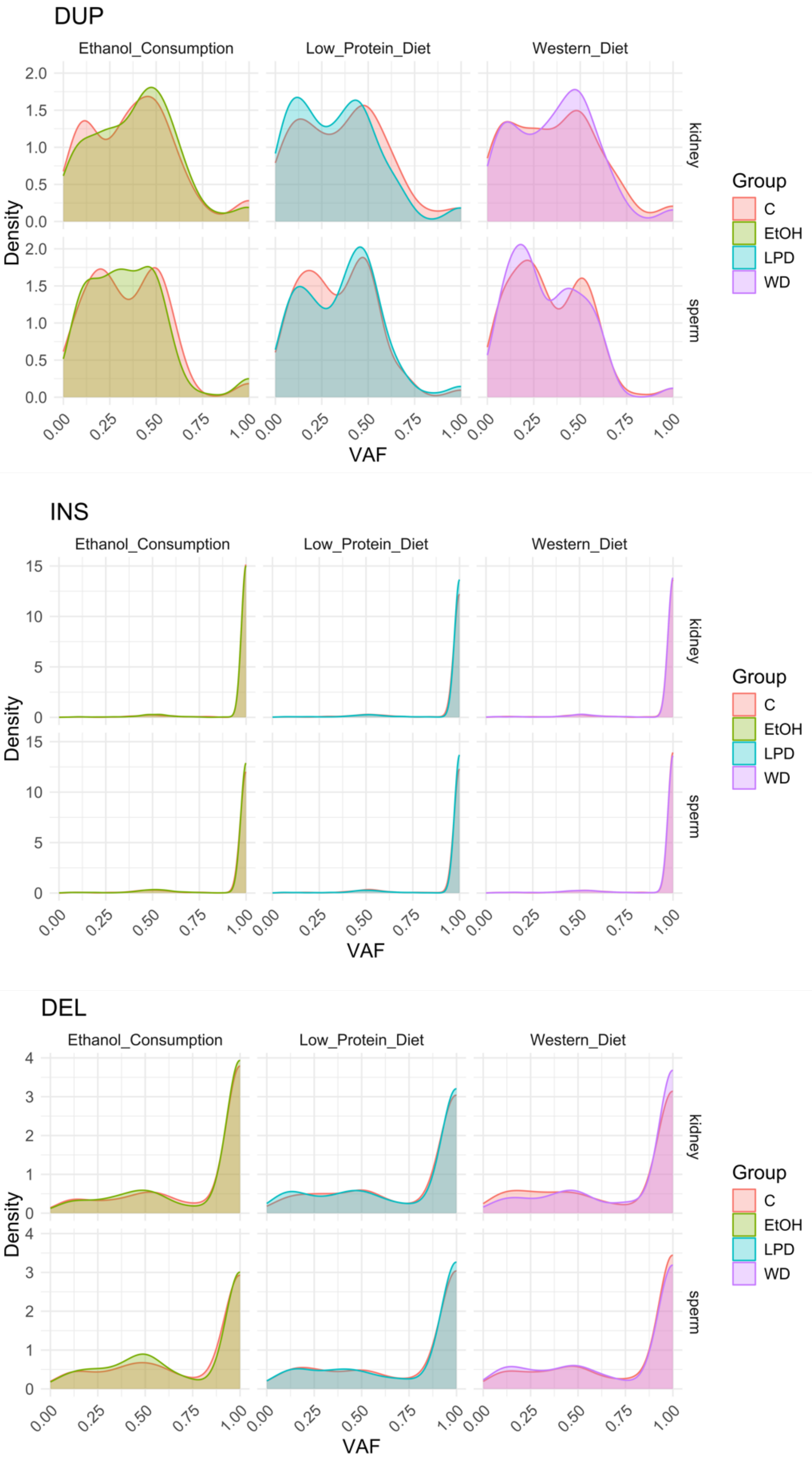


**Fig. S5.** Comparisons of VAF distributions between groups and by SV type.

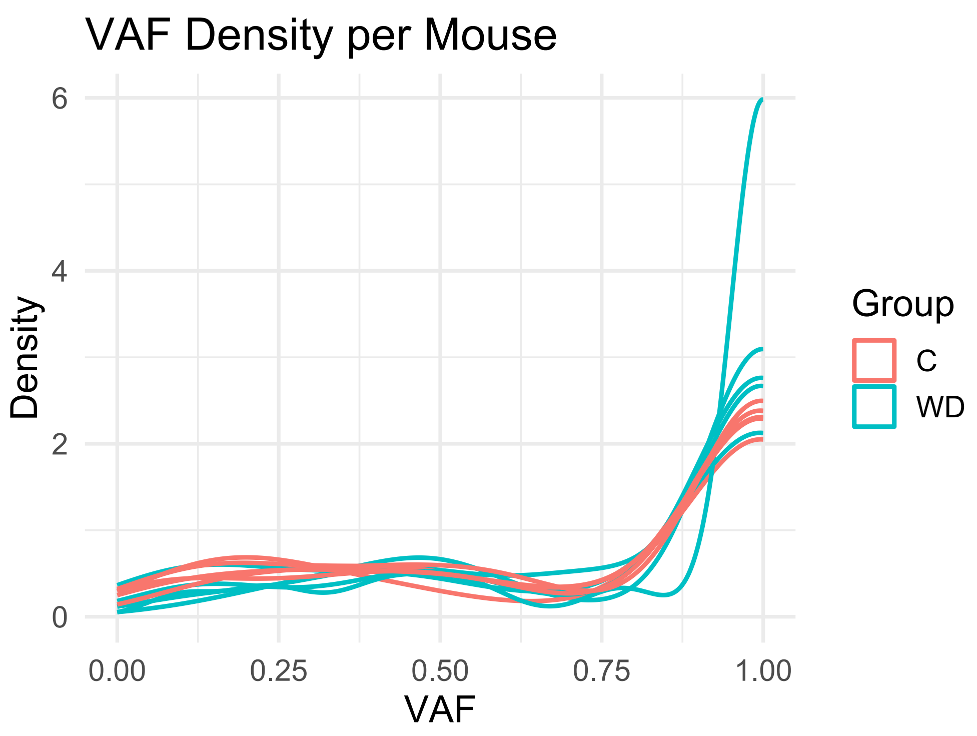


**Fig. S6.** Comparison of VAF distribution of deletions between kidney samples of individual mice in the Western Diet experiment.

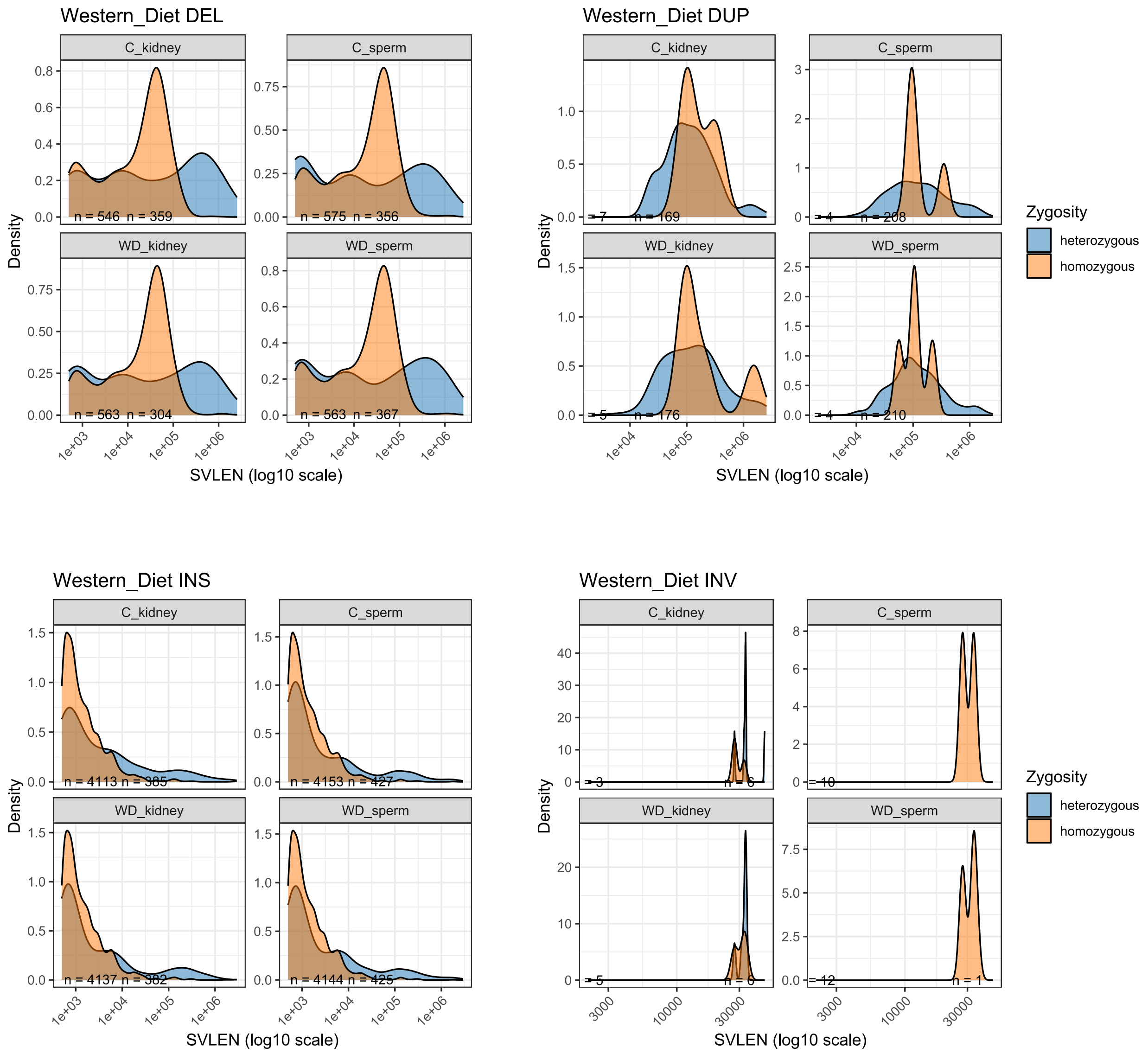


**Fig. S7.** Distribution of homozygous and heterozygous SVs in the Western Diet experiment

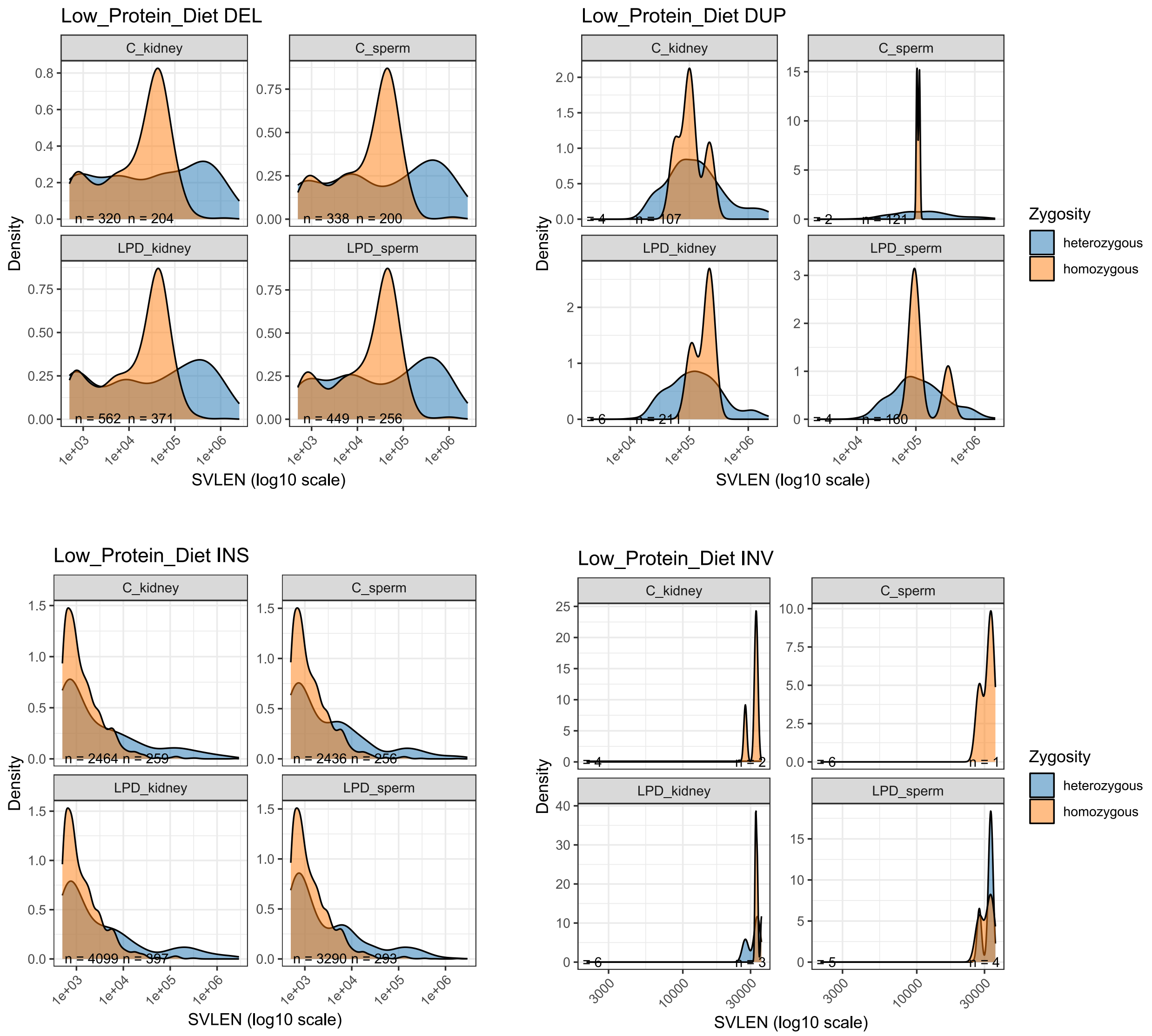


**Fig. S8.** Distribution of homozygous and heterozygous SVs in the low-protein diet experiment

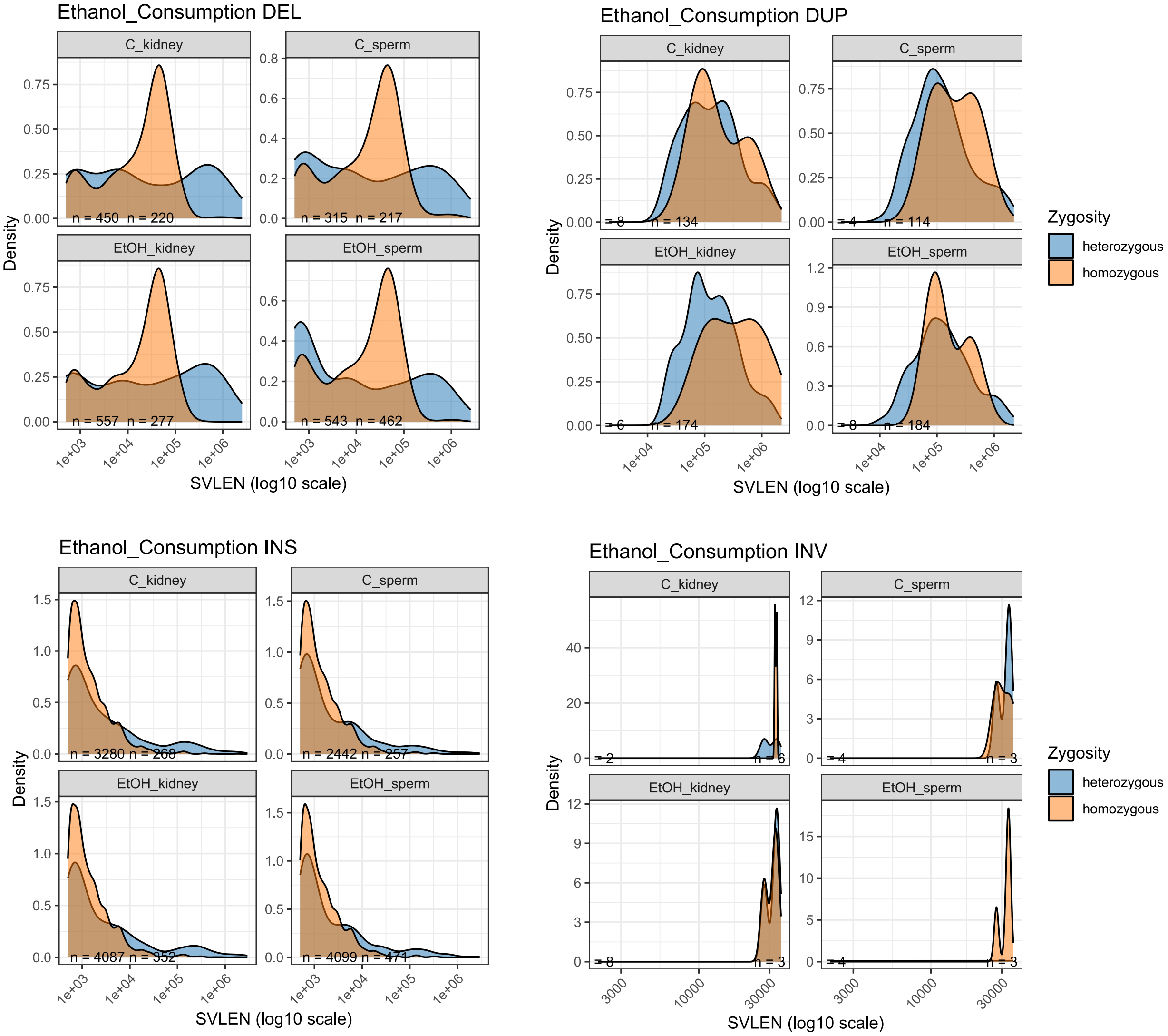


**Fig. S9.** Distribution of homozygous and heterozygous SVs in the ethanol consumption experiment

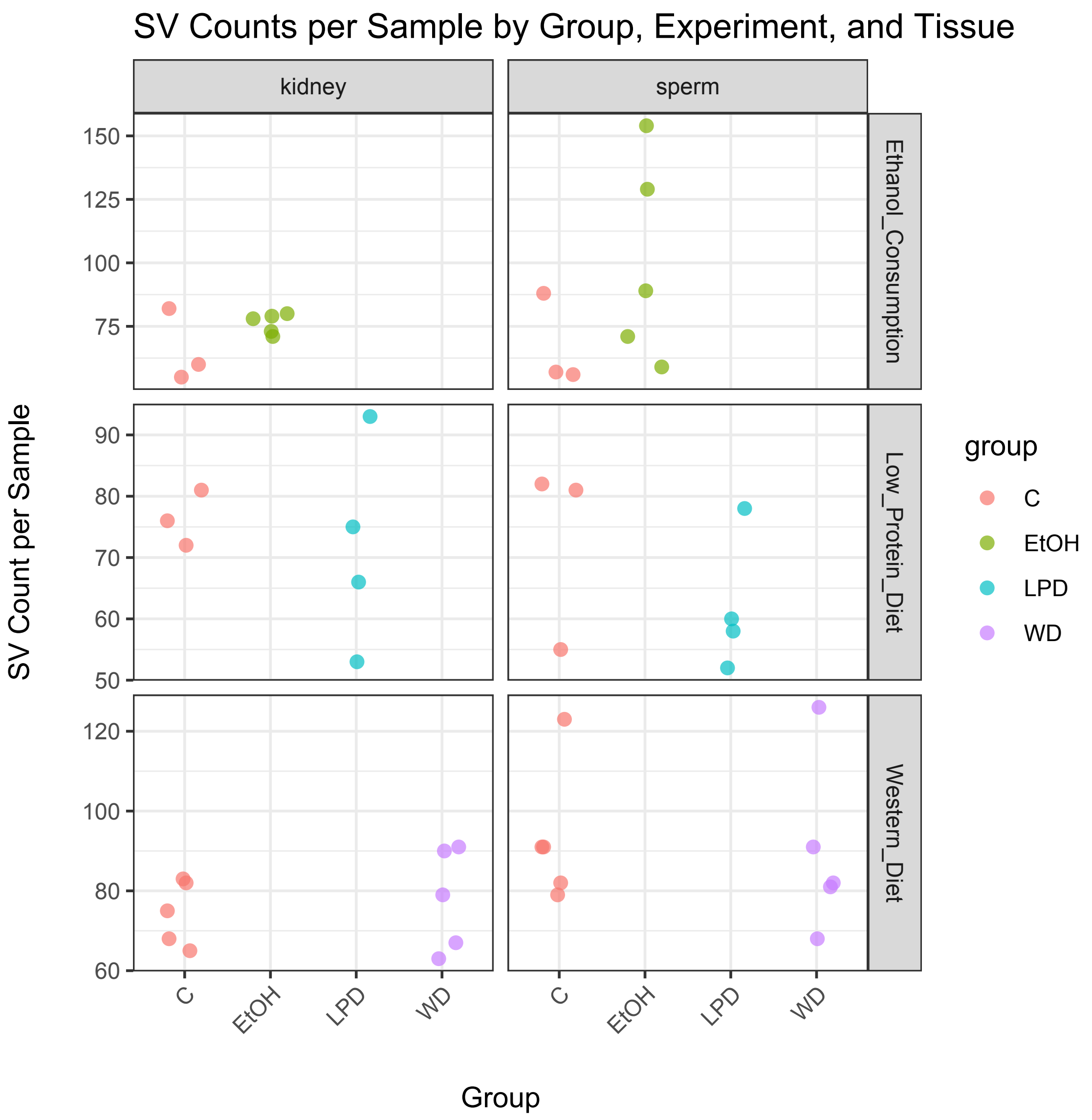


**Fig. S10.** Number of detected de novo SVs per mouse

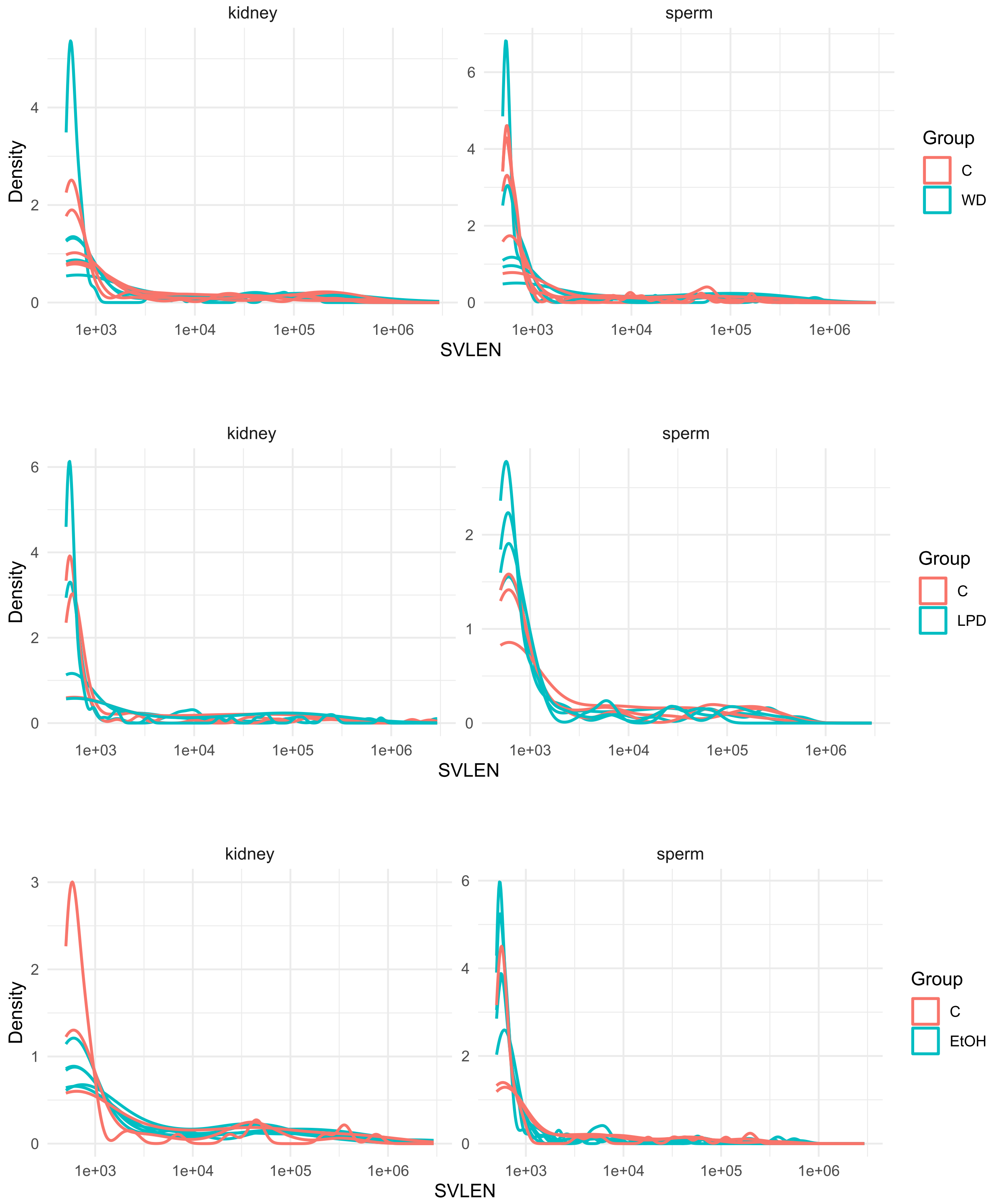


**Fig. S11.** Distribution of lengths of tissue-specific de novo SVs.

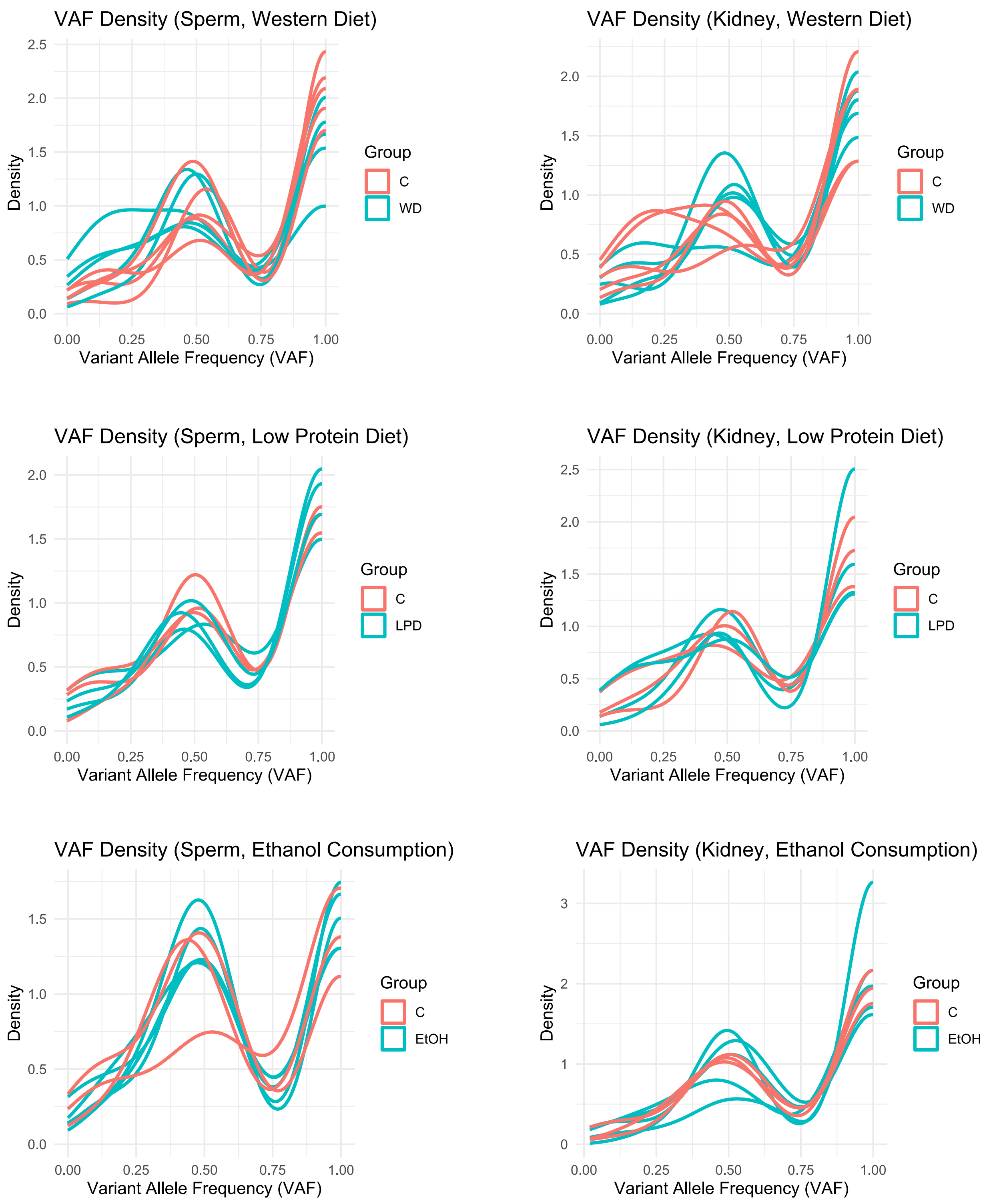


**Fig. S12.** Distribution of variant allele fraction of tissue-specific de novo SVs.
